# An automated respiratory data pipeline for waveform characteristic analysis

**DOI:** 10.1101/2022.12.02.518741

**Authors:** Savannah Lusk, Christopher S. Ward, Andersen Chang, Avery Twitchell-Heyne, Shaun Fattig, Genevera Allen, Joanna Jankowsky, Russell Ray

## Abstract

Comprehensive and accurate analysis of respiratory and metabolic data is crucial to modelling congenital, pathogenic, and degenerative diseases converging on autonomic control failure. A lack of tools for high-throughput analysis of respiratory datasets remains a major challenge. We present Breathe Easy, a novel open-source pipeline for processing raw recordings and associated metadata into operative outcomes, publication-worthy graphs, and robust statistical analyses including QQ and residual plots for assumption queries and data transformations. This pipeline uses a facile graphical user interface for uploading data files, setting waveform feature thresholds, and defining experimental variables. Breathe Easy was validated against manual selection by experts, which represents the current standard in the field. We demonstrate Breathe Easy’s utility by examining a 2-year longitudinal study of an Alzheimer’s Disease mouse model to assess contributions of forebrain pathology in disordered breathing. Whole body plethysmography has become an important experimental outcome measure for a variety of diseases with primary and secondary respiratory indications. Respiratory dysfunction, while not an initial symptom in many of these disorders, often drives disability or death in patient outcomes. Breathe Easy provides an open-source respiratory analysis tool for all respiratory datasets and represents a necessary improvement upon current analytical methods in the field.

## Introduction

Whole-body plethysmography (WBP) in rodent models is an invaluable tool for investigating the etiology and progression of many diseases including congenital respiratory disorders such as Sudden Infant Death Syndrome, contagious respiratory viruses such as SARS-CoV-2, respiratory dysfunction derived from spinal cord and brain injuries, and loss of airway protection and respiratory failure in neurodegenerative disorders such as Alzheimer’s Disease. Elaborated WBP systems can measure the volume and composition of air that is inhaled and exhaled per breath offering novel and critical insights into disease etiology and progression. This technique has become integral in the analysis of respiratory outcomes leading to an average of over 600 articles every year published using this method for the last 20 years. Despite the importance of this technique, its application remains limited as few, if any, comprehensive, open-source tools exist for respiratory waveform analysis, leaving manual data annotation as the dominant approach. To aid in the consistent analysis of respiratory waveform and metabolic data, and to make possible high-throughput and immensely sized experimental designs, we have developed the Breathe Easy software package.

The analysis of plethysmography data consists of two components: 1) the conversion of collected waveforms into physiological measures, e.g., respiratory frequency and tidal volume, and 2) the statistical assessment and graphical display of those measures and their associative relationships with independent variables. Labor-intensive, time-intensive, and expertise-dependent manual selection of data remains the most common approach^1, 2^, which results in a significant bottleneck. Current annotation methods limit reproducibility within and between labs (Supplemental Table 1); is subject to observer bias (i.e., where humans tend to select for particularly slow, even breaths), especially when identifying and excluding movement artifacts; and typically includes only a tiny sample of all available data (usually only examining 30 seconds – 2 minutes of breathing per experimental condition^1–3^). Analysis of the selected data can be similarly encumbered by a heavy reliance on more manual approaches that involve copying and pasting manually-selected data into inherited spreadsheets with default formulae^2, 3^. Each component of the analysis is manually driven, which makes analysis of data from even the simplest of experiments exceedingly time consuming. The few available commercial solutions that enable partially automated analysis are 1) expensive; 2) closed source; 3) remain limited in their overall capabilities; and 4) often require purchase of a commercial plethysmograph system for full analytical options, which precludes analysis of data collected on other systems and the use of custom WBP systems (Supplemental Table 2). These limitations further constrain the complexity and magnitude of respiratory studies and dissuade multivariate experimental designs examining interactions across multiple variables such as genotype, drug, pathogen, injury, physiological challenges (e.g., hypercapnic, hypoxic, or room air breathing), sex, and age (in the case of longitudinal studies). Thus, approaches are lacking that enable the customized evaluation of large or complex (multivariate) datasets for cardiorespiratory and metabolic physiology studies.

Here we present Breathe Easy, a novel, modular pipeline that can be run unsupervised in a continuous fashion over large datasets or more flexibly for highly customized analyses, thereby overcoming the above limitations. Breathe Easy consists of three modules: 1) a graphical user interface for rapid and facile loading of data and customization of parameters surrounding waveform feature extraction, desired statistical analysis, and graphical displays; 2) a breath extraction module to segment and quantify pertinent respiratory variables using pressure, temperature, and gas concentration waveforms filtered by exclusion criteria; and 3) an analysis and graphing module that generates publication-worthy graphs and statistics accompanied by QQ and residual plots to gauge adherence to assumptions. We first validated Breathe Easy to show that it outperforms commonly used manual annotation techniques. We then used Breathe Easy to analyze a large, longitudinal respiratory dataset collected over two years (more than 600 WBP sessions) using a genetic model of Alzheimer’s Disease (AD) and demonstrated that our software robustly handles a multitude of waveform anomalies and can readily manage large terabyte-scale respiratory data sets.

## Results

### Overview of software

Figure 1 provides an overview of our new modular, comprehensive, and automated respiratory analysis software pipeline, Breathe Easy, which is flexibly designed for use both by novices and experts. More detailed descriptions of each component of Breathe Easy can be found in the Methods section. To use Breathe Easy, we strongly recommend new users consult our User Manual website, which can be accessed through the GitHub repository linked to this manuscript. Briefly, the data and user-defined inputs for the pipeline are managed through the main Graphical User Interface (GUI), increasing accessibility to novices both in respiratory waveform analysis and coding (Fig. 1A). The overarching workflow outlined in Figure 1B for a typical multivariate respiratory experiment is managed through a series of subGUIs (Fig. 1C) that allow the user to define inclusion and exclusion criteria for waveform features, parameters for the experimental design, and options for statistical analysis and graphical output. Screenshots of each subGUI are shown in Supplemental Figure 1.

**Figure 1.**
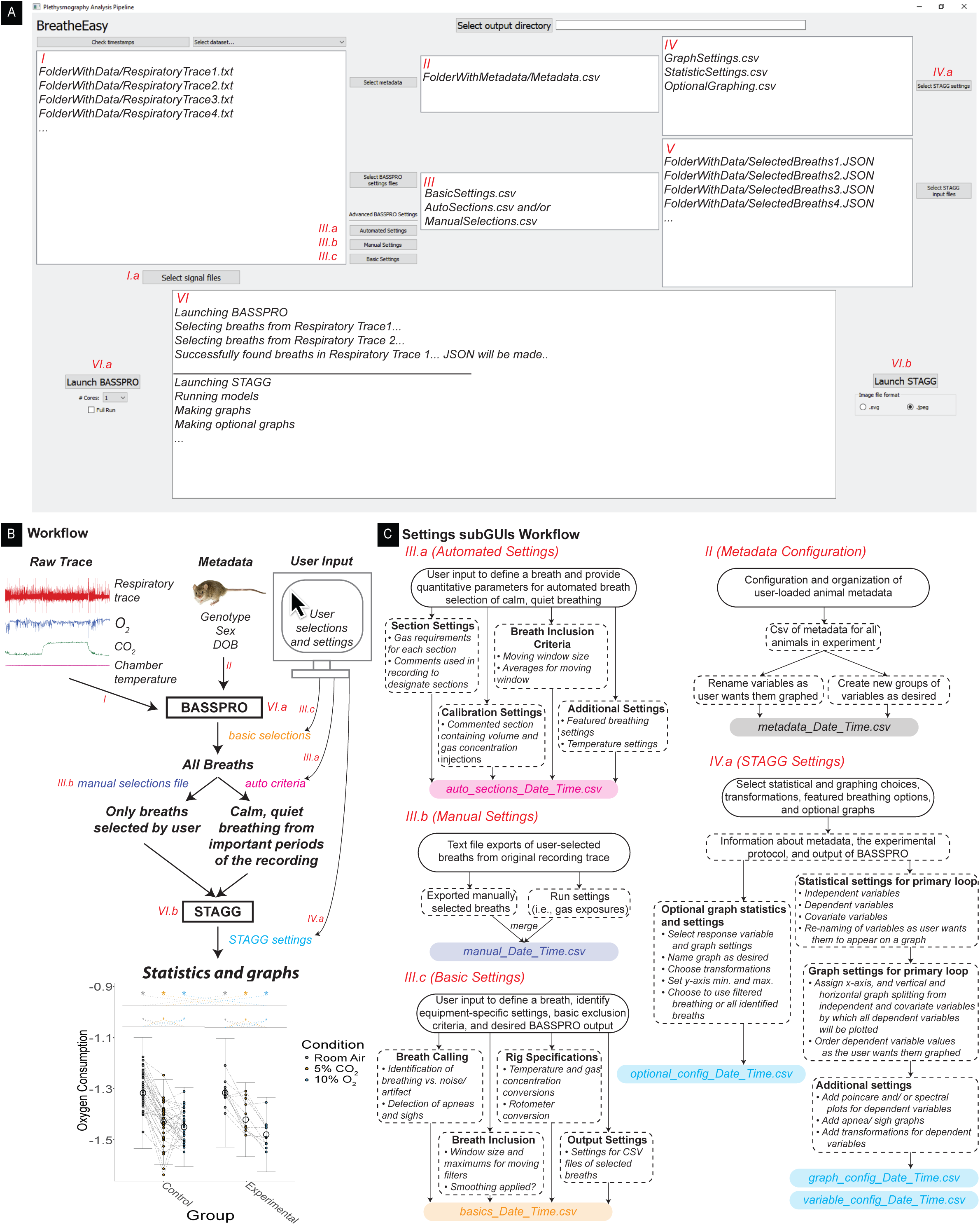
Overview of Breathe Easy components and workflow. A) Screenshot of the main GUI interface where the red numbering corresponds to displays shown in B-C. Text shown in white boxes are not what the user will see in the GUI, rather an example of what will populate those areas when using the GUI. B) A walk through of the full workflow of Breathe Easy with reference to each component in the GUI. C) Workflows and outputs of all of our subGUI interfaces.

The raw plethysmography signal files, experimental metadata files, and user-defined filtering parameters (i.e., configuration files generated by the GUI based on user selections) are passed to the Breathing Analysis Selection and Segmentation for Plethysmography and Respiratory Observations (BASSPRO) module. The configuration files inform BASSPRO how the user defines a breath and exclusion criteria for artifacts like animal movement or flow obstructions. The application of these criteria is demonstrated in an example trace in Figure 2A. BASSPRO records instantaneous parameters for each identified breath that is selected for inclusion and compiles the output in a single .json file for each recording.

**Figure 2.**
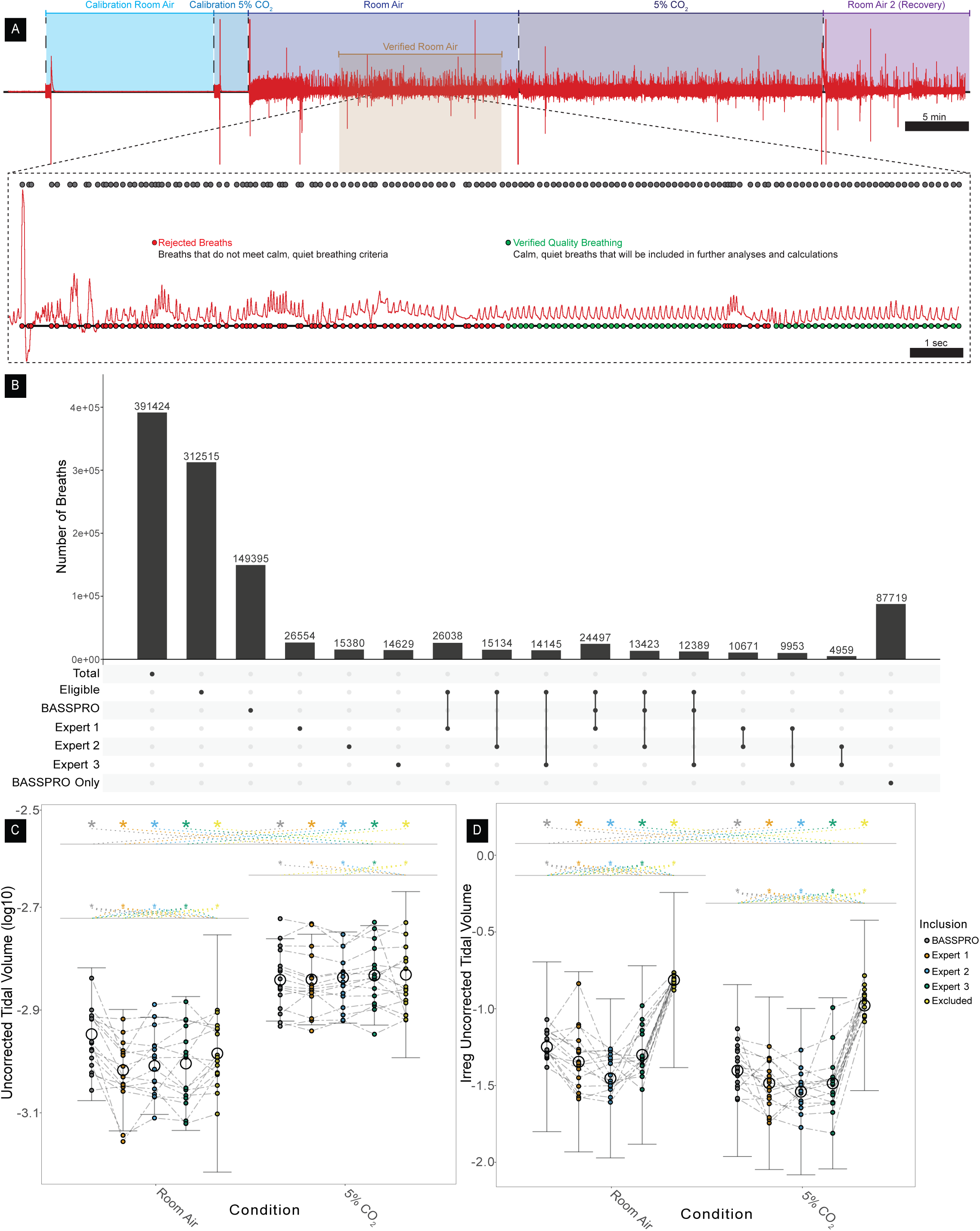
Demonstration of breath selection validation where A) shows selection process behind the scenes of BASSPRO where gray overhead dots show all breaths identified and the red and green dots below the trace show the decision to include (green) or exclude (red) each breath and B) shows a comparison of breaths selected by BASSPRO to those manually selected by 3 experts from the same 16 files. C-D are uncorrected tidal volume and irregularity score of uncorrected tidal volume for breaths selected by BASSPRO vs manual selections by 3 experts. Error bars = standard error mean, * = p<0.05.

Following quality control, waveform filtering, and feature detection by BASSPRO, the data is sent to the STAtistics and Graph Generator (STAGG) module where statistics and graphs are generated based on user-defined settings. The user selects independent, dependent, and covariate variables for statistics; assignments for graphing respiratory variables; instructions for graphs other than respiratory variables selected by the user (e.g., body weight, body temperature, behavioral experiment outcomes, etc.); or different graphing settings for respiratory variables.

A successful run of Breathe Easy generates publication-ready figures with statistical significance marks added and tables of quantitative data representing the graphical outputs and showing statistical testing results. The output of this pipeline and required efforts to run the program represent a significantly improved and streamlined pipeline for plethysmography data analysis.

### Validation

To validate the automated selection of breaths by BASSPRO, we compared the breaths selected by BASSPRO to those selected manually by experts in 16 representative traces from the 14 months timepoint in our Alzheimer’s dataset. Three experts (as defined by a record of peer-reviewed publication in respiratory waveform analysis^4–6^) were engaged to select periods of calm, quiet breathing from regions of interest (i.e., stable gas concentrations for room air and challenge gas exposure) for analysis and comparison. The same files were then run through Breathe Easy using default parameters. Out of 391,424 total breaths identified in the 16 traces, 312,515 breaths were from desired sections of the recording (e.g., animal habituated and room air or fully equilibrated 5% CO_2_). Each expert selected, on average, 18,854.3 ± 6,678.7 breaths whereas BASSPRO selected 149,395 breaths from those same 16 files (Fig. 2B). 88.1% ± 3.8% of breaths selected by each of the experts were also selected by BASSPRO, showing high overlap between the two selection methods. In contrast, 11.9% ± 3.8% of breaths selected by experts were rejected by BASSPRO, where ∼3% of this difference is due to experts selecting outside of the desired periods of interest, e.g., 6 minutes before the end of the condition instead of only selecting from the last 5 minutes. Thus, of breaths selected by experts as calm, quiet breathing from the correct periods of interest, less than 9% of breaths selected by experts were rejected by BASSPRO for not meeting quality standards.

There is concordance between BASSPRO and the experts and high discordance between excluded breaths and those included by BASSPRO and experts when considering the average of basic breathing variables calculated from breath selections as shown in Supplemental Table 3 including breath cycle duration (A), ventilatory frequency (B), and uncorrected tidal volume (C, Fig. 2C). There is also a statistically significant difference in the irregularity scores for basic variables when comparing breaths selected by BASSPRO to those excluded by BASSPRO as shown for uncorrected tidal volume (Fig. 2D). We also see concordance between BASSPRO and the experts with the averages of refined respiratory variables as shown in Supplemental Table 3 including peak inspiratory flow (D), peak expiratory flow (E), ventilatory equivalents of oxygen (F), corrected tidal volume (G), weight normalized ventilation (H), and oxygen consumption (I).

### Biological Application

AD is a chronic, neurodegenerative disease with only palliative treatments currently available. As neurodegeneration worsens, key bodily functions including respiration begin to fail, which directly or indirectly leads to death. Sleep-disordered breathing (SDB) and aspiration pneumonia show strong associations with AD progression. There is a significantly increased prevalence of SDB in patients with dementia compared to the general population (71% vs. 55.7%)^7–9^, and the E4 allele of apolipoprotein E, which is the strongest genetic risk factor for AD, is more highly expressed in patients with sleep apnea^10^. Continuous positive airway pressure for treatment of SDB in AD patients may improve symptoms^11^. Additionally, a leading cause of morbidity in AD and other neurodegenerative pathologies is aspiration pneumonia, which is hypothesized to result from disruption and degeneration of protective brainstem respiratory and swallowing networks^12–19^. The summation of the evidence suggests that poor breathing function may play a role in AD onset, progression, and severity, and that AD progression may create a mutually destructive feedforward loop that disrupts respiratory networks.

To gain insight into respiratory function during AD progression, we used WBP on an inducible amyloid precursor protein (APP) forebrain mouse model^20^ to characterize metabolic and respiratory function over nearly two years. Along with room air breathing (21% O_2_/79% N_2_), hypercapnic (5% CO_2_/21% O_2_/74% N_2_) and hypoxic (10% O_2_/90% N_2_) respiratory homeostatic chemosensory functions were assayed. We focused on chemosensory respiratory reflexes as these neurological reflexes are often perturbed in SDB and other respiratory pathophysiologies.

A total of 633 recordings were completed in the 2-year longitudinal Alzheimer’s study following 55 mice across experimental and control genotypes. Beginning approximately 5 months after cessation of Doxycycline treatment, immediately following the first appearance of amyloid plaques, mice were recorded about every 2 months for 19 months. Timepoints reported in figures and text refer to total time off Doxycycline treatment. Not all mice met inclusion criteria for all timepoints; some died of age-related complications or recordings were not of sufficient quality. Because our analyses are completed on a breath-by-breath level rather than average values for each animal, we were able to include data from at least one period of the recording for 44 animals, resulting in 302 recordings for hypercapnic ventilatory response and 301 recordings for hypoxic ventilatory response, which are included in the data shown in Figures 3-6.

**Figure 3.**
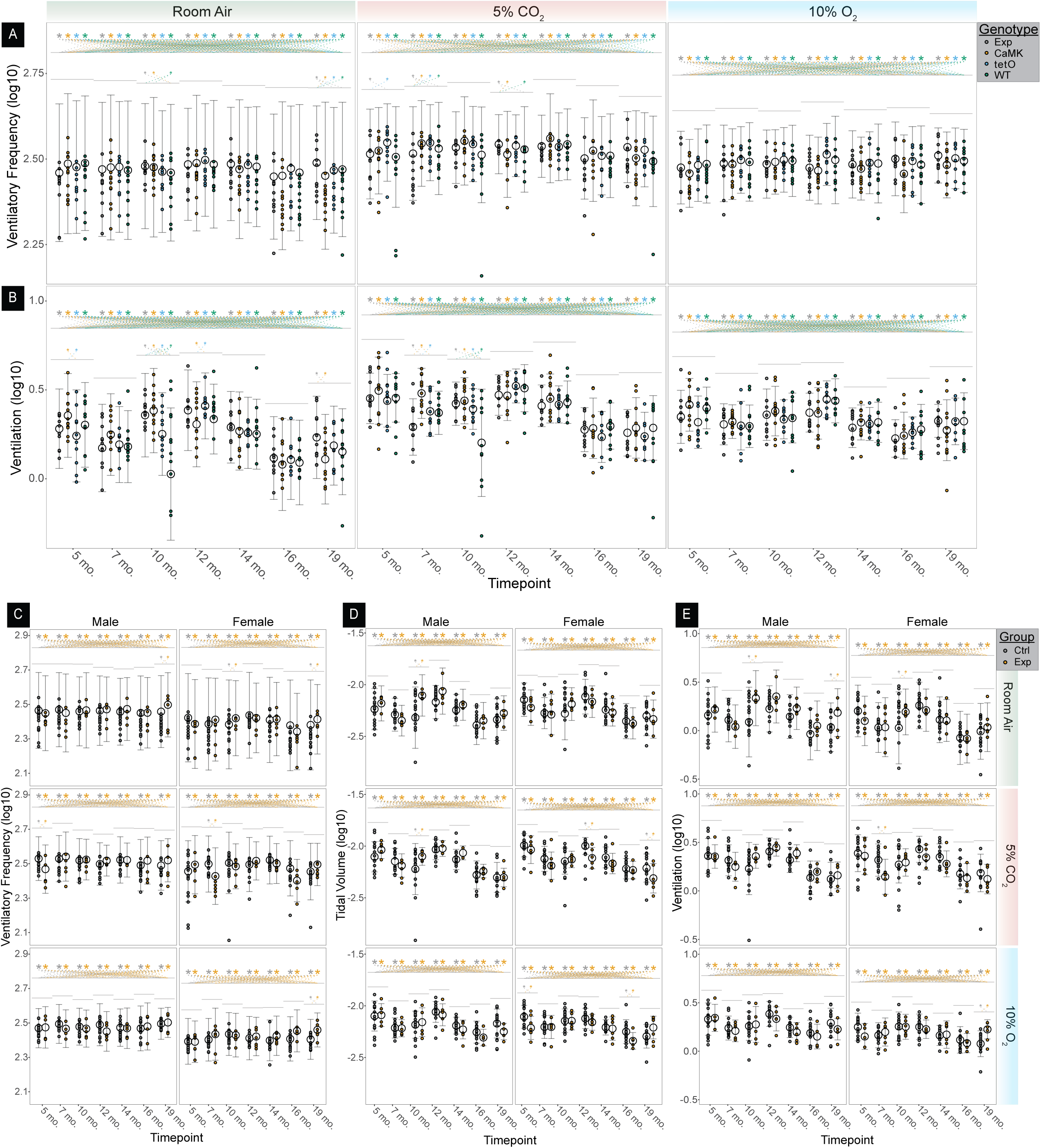
Respiratory outcomes from our 2-year longitudinal Alzheimer’s study. A) Ventilatory frequen- cy and B) Ventilation plotted by timepoint, genotype, and condition with sex considered in the statisti- cal model as a covariate. C) Ventilatory frequency, D) Tidal volume, and E) Ventilation plotted by timepoint, group, condition, and sex. Error bars = standard error mean, * = p<0.05.

No statistically significant trend in respiratory variables was observed in the Alzheimer’s group compared to age-matched controls over time. While some timepoints showed significant differences between experimental and control animals or within the same group over time, these differences presented as random with no obvious biological cause. A number of the significant differences were due to significant variability in one or more of the control groups: ventilatory frequency (*V_F_*) at 10 months in room air, and ventilation (*V̇_E_*) at 5 months in room air, 10 months in room air, 12 months in room air, 19 months in room air, 7 months in 5% CO_2_, and 10 months in 5% CO_2_ (Fig. 3A-B, Supplemental Table 4). There are also cases where just one or two of the control groups differ from the experimental group: *V_F_* at 5 months in 5% CO_2_ and 12 months in 5% CO_2_ (Fig. 3A, Supplemental Table 4).

Breathe Easy is designed so the user can easily assess the effects of grouped or individual levels within independent variables across multiple variables in an experiment to uncover key factors involved in differential outcomes. For example, in the case of *V_F_* at 5 months in 5% CO_2_, we can see that this difference is driven completely by the males, whereas the difference at 12 months in 5% CO_2_ no longer holds up when the control groups are combined, and sex is added as a factor (Fig. 3C). Another example of deconvoluting outcomes can be found by examining individual levels for sex and genotype variables in *V_F_* and *V̇_E_* at 7 months in 5% CO_2_ where it becomes clear that the difference is driven by a change in *V_F_*, but not tidal volume, in the females leading to a significant difference in *V̇_E_* (Fig. 3C-E). However, this difference in *V̇_E_* seems chiefly caused by the CaMKII-tTA control group when the control groups are split, and sex is removed as a factor. This is in line with previous findings indicating phenotypes with the CaMKII-tTA driver to be commonplace^21^.

Upon further investigation into metabolism, we see similar trends for seemingly random significant differences across a number of comparisons that do not have an obvious biological cause; these significant results are also not sustained when independent variables are grouped differently (Fig. 4A,C). However, we do note significant differences at 7 months as we saw with *V̇_E_* and *V_F_*. In 5% CO_2_ there is a significant difference between experimental and control animals in ventilatory equivalents of oxygen (*V̇_E_*/*V̇_O_2__*), but not oxygen consumption (*V̇_O_2__*) (Fig. 4A-B). This difference persists in both males and females for *V̇_E_*/*V̇_O_2__* whereas no difference is noted in *V̇_O_2__* (Fig. 4C-D). This suggests, combined with the data shown in Fig. 3B,E, that this difference is driven by a change in *V̇_E_* and not *V̇_O_2__*. Further, although the difference in *V̇_E_* was not significant in males (Fig. 3E), there is significance in males for *V̇_E_*/*V̇_O_2__*. This suggests that a trend in *V̇_E_* for males may be present, but not significant, and becomes clear when normalized to metabolic demand.

**Figure 4.**
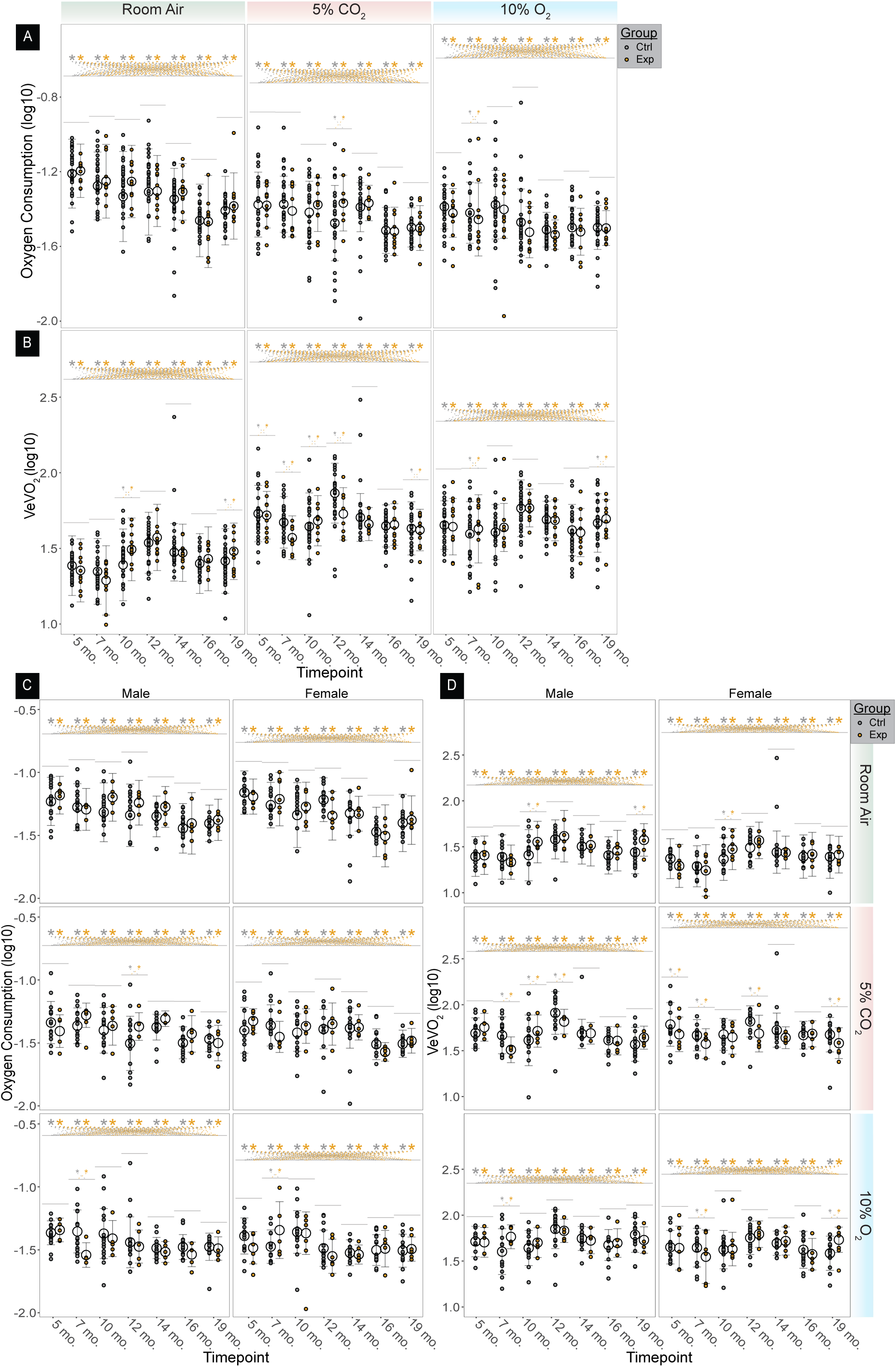
Respiratory and metabol- ic outcomes from our 2-year longi- tudinal Alzheimer’s study. A) Oxygen consumption and B) Ventilatory equivalents of oxygen plotted by timepoint, genotype, and condition with sex considered in the statistical model as a covariate. C) Oxygen consumption and D) Ventilatory equivalents of oxygen plotted by timepoint, group, condi- tion, and sex. Error bars = standard error mean, * = p<0.05.

On the other hand, in 10% O_2_, we see no differences in *V̇_E_* at 7 months (Fig. 3B,E), but we do see significant differences in *V̇_O_2__* and *V̇_E_*/*V̇_O_2__* even when the data is split by sex (Fig. 4), which suggests this difference is driven by a change in *V̇_O_2__*. Interestingly, we see a bimodal distribution of the *V̇_O_2__* response where males show a decrease and females show an increase. Despite this differential response, the change in *V̇_E_*/*V̇_O_2__* between control and experimental animals still holds when sex is combined (Fig. 4B).

We also report the featured breathing phenotypes for apneas and sighs in this study where we do not observe any meaningful changes in sigh rate that would be indicative of a phenotype driven by this Alzheimer’s model (Fig. 5B,D). For apnea rate, there are a number of significant differences between controls and experimental animals in females under 5% CO_2_ when split by sex (Fig. 5A), but none of those changes remain significant when sex is removed (Fig. 5C).

**Figure 5.**
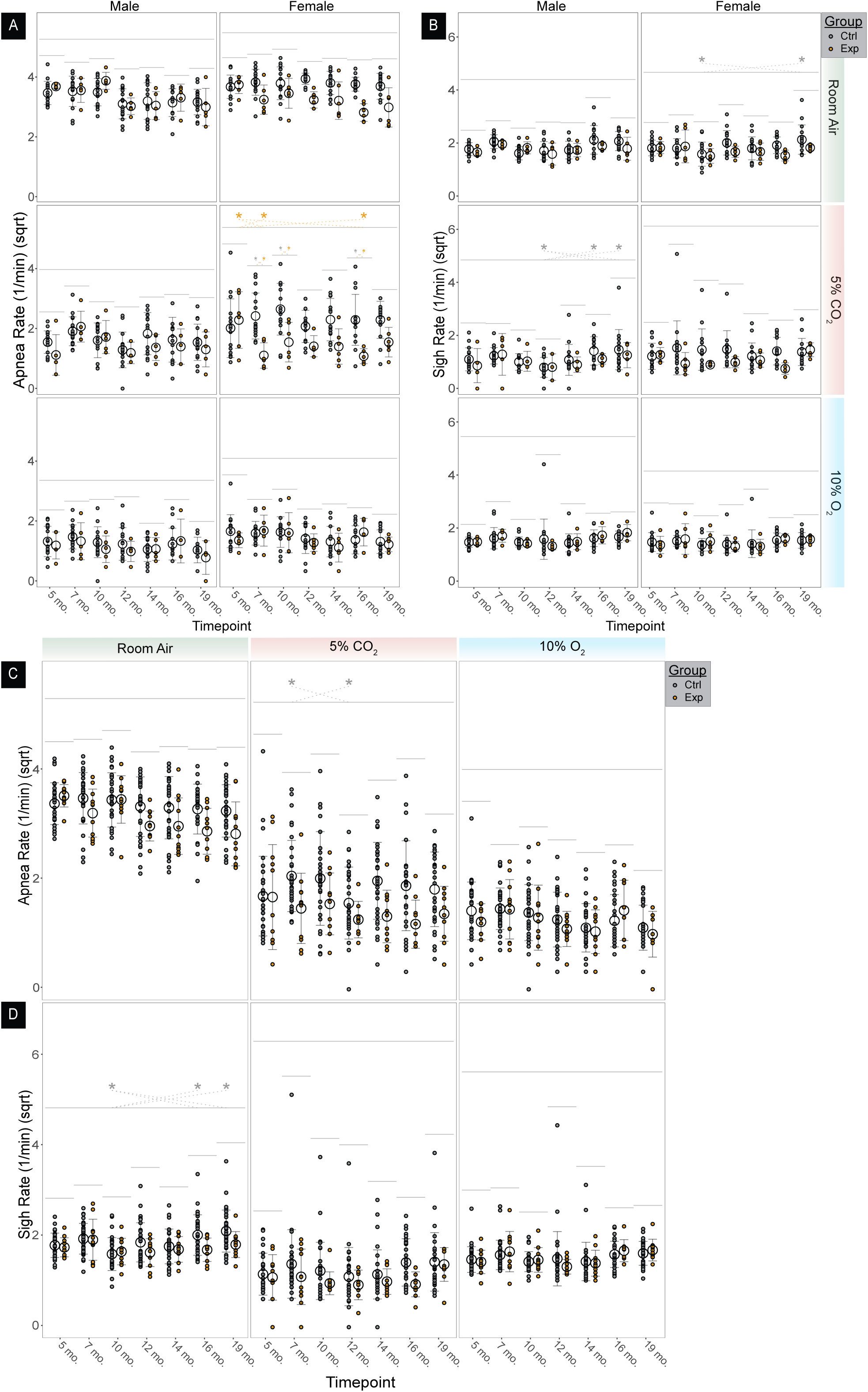
Apnea and sigh outcomes from our 2-year longitudinal Alzheimer’s study. A) Apnea and B) Sigh plotted by timepoint, group, condition, and sex. C) Apnea and D) Sigh plotted by timepoint, group, and condition with sex considered in the model as a covariate. Error bars = standard error mean, * = p<0.05.

There is also a difference in experimental animals between 5 and 7 months, and 5 and 16 months in 5% CO_2_ (Fig. 5A), however this difference is also gone when sex is removed and instead we see a difference in control animals between 7 and 12 months in 5% CO_2_ (Fig. 5C).

Additionally, we can easily test for contributions of known related variables like weight and age. No significant difference between experimental and control animals was seen in the number of animals included or weight at the time of their recordings (Fig. 6A, Supplemental Table 5) aside from the expected significant weight gain with age. There was a significant difference in age between experimental and control animals at 12 months (E=experimental average±standard error, C=control average±standard error, p-value) (E= 400.1±14.0, C= 393.6±12.4, p=0.0073), 14 months (E= 459.3±10.8, C= 452.8±12.3, p=0.021), 16 months (E= 518.1±8.67, C= 510.6±9.52, p=0.022), and 19 months (E= 601.2±6.46, C= 593.3±9.4, p=0.022) (Fig. 6B).

**Figure 6.**
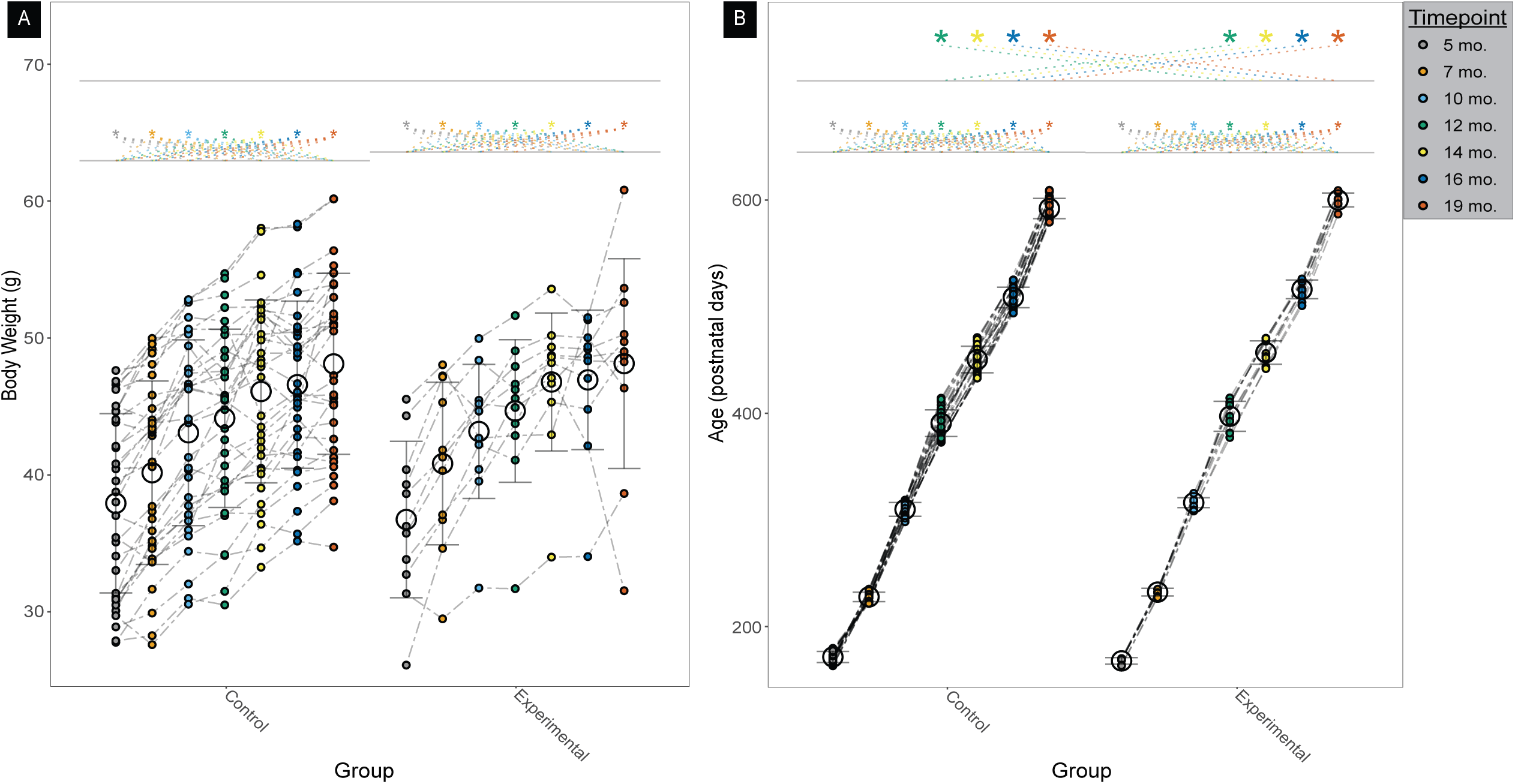
A) Body weight and B) Age of animals used in our longitudinal Alzheimer’s study plotted by timepoint and group. Error bars = standard error mean, * = p<0.05.

Although the ages are significantly different, we expect that this significance is derived from the order by which the animals were run through our experimental paradigm. Unfortunately, it was not considered that the genotypes and groups should be randomly recorded at each of the timepoints, so the animals were run by group in the same order for each timepoint. This led to a significant, but not biologically relevant difference in age at the final timepoints. This hypothesis is supported by the consistent decrease in variation in age for both groups as age increases.

## Discussion

Here we present a novel software platform that offers a significant improvement upon current analysis methods for WBP data. While commercial options for partially automated analyses are available, they are closed source and offer little flexibility for breath identification and filtering, and subsequent analysis (Supplemental Table 2). While several approaches to analyzing differing aspects of rodent respiratory data have been published in one form or another^22–26^, no available software suite, to our knowledge, has the capability of filtering for the typical standard of calm, quiet breathing in rodent plethysmography experiments, i.e., automatically filters movement and other waveform artifacts. In addition to programmatic benefits outlined in Supplemental Table 2, Breathe Easy is open source, free to academic researchers, and offers a novel, raw data-to-figure pipeline.

Conventional approaches to identification and selection of breaths for analysis fall into two extremes: manual selections or inclusion of all breaths. The BASSPRO module of Breathe Easy provides its users with the ability to identify all breaths within a recording and provide a filtered selection of breaths that generally agree with expert selections of “calm, quiet” breathing. The automated selections provided by BASSPRO have the advantage of being impartial and tractably defined by user-adjustable settings that can easily be shared across multiple users, eliminating the problem of inter-rater reliability when analyzing large datasets. Additionally, BASSPRO avoids using the breathing rate of individual breaths as a selection filter. Instead, it uses a sliding window that incorporates neighboring breaths to exclude breaths proximal to very high frequency breathing, such as bouts of sniffing, while permitting inclusion of a larger range of likely “calm, quiet” breaths throughout the full duration of the recording. This range is lost with manual selection as experimenters are prone to selecting the slowest and steadiest breathing intervals. BASSPRO also integrates information across other data channels to aid in defining the filtering criteria for automated selections. BASSPRO additionally has the capability of providing indirect calorimetry respiratory outcomes, which are essential as a means of controlling for the effects of metabolic demand on breathing outcomes. Limitations of the current implementation relate to the initial candidate breath identification settings as the same settings are used for candidate breath detection for the full recording and for all recordings analyzed as a batch. This may require a user to refine and repeat their analysis in cases where marked differences in respiratory flow are present across animals, or across portions of the recording.

Breathe Easy’s STAGG module is also unique in the application of appropriate statistical testing for respiratory datasets. While many articles in the field apply ANOVA or t-tests, these tests are mostly inappropriate for the structure of most data collected in plethysmography experiments, which often involve repeated measurements on the same subjects, particularly for chemosensory assays, and naturally involve multiple variables that influence breathing but are not always considered, e.g., age, sex, and experimenter. Our application of linear mixed effects models (LMEM) with Tukey post hoc analysis represents a needed standardization for analysis in the field and, arguably, a significant improvement in statistical analysis, as the model accounts for repeated measures and accommodates more complex experimental designs.

Several assumptions are made in the application of mixed effects models that should be addressed by the researcher before interpreting results. We use a standard LMEM model for all dependent variables, in which it is assumed that the random effect coefficients and the overall model residuals follow normal distributions. Cases where the residuals are not independent and identically distributed indicate potential invalidity of the model. While strict adherence to assumptions with an LMEM is not always required^27^, we do provide output to address these assumptions. QQ and residual plots are generated for each dependent variable. Based on these, the user can choose to include models with transformed dependent variables to address adherence to these assumptions. Transformations that are currently available include log_10_, natural log, and square root.

While the LMEM is widely used to analyze data in biological contexts, the standard model we use in STAGG for all designated dependent variables has some potential limitations. One issue is that the standard LMEM is only applicable for continuous dependent variables. If a user wanted to analyze categorical or discrete dependent variables, this would require the usage of a generalized linear mixed effects model, which is currently not implemented in STAGG. In cases of optional graphs where this would be appropriate, an ANOVA is used instead of the primary LMEM model. Additionally, the standard LMEM model does not account for any possible dependencies between subjects or any correlations between observations outside of what is accounted for in the random effects. For example, there could be autocorrelation effects from a single session or possible additional repeated measures effects from the experimental design, which the current implementation does not handle. However, it is possible for the user to run STAGG to test these dependencies by selecting rig, experimental date, time of day, etc. from the imported metadata as dependent variables to test for significant contributions.

Breathe Easy was validated against the typical standard of manual selection of recordings (Fig. 2). The concordance between BASSPRO selected breaths and all three experts demonstrate the ability of the software to select breathing that an expert would deem calm, quiet breathing.

Furthermore, as sample size increases, we know the accuracy of estimated mean becomes closer to the true mean. We see a significant difference between breaths selected by BASSPRO and breaths excluded by BASSPRO; therefore, the accuracy of BASSPRO is not just due to an increase in sample size and is instead due to the quality of selected breaths. We also see significant variation between experts, which is not uncommon in respiratory experiments where reproducibility both in values and units reported is questionable (Supplemental Table 1). This problem can easily be solved by adoption of Breathe Easy across labs performing WBP experiments to ensure consistent data analysis.

The high level of variance in agreement between experts, and between experts and BASSPRO- selected breaths is encouraging as it supports and provides evidence for the argument of increased consistency when using Breathe Easy compared to manual selection. It is not uncommon in the field of respiratory studies for experiments to have unreliable reproducibility.

These data show that even with a blinded group of experts there is variation in the manual selections that lead to significantly different results. Furthermore, while BASSPRO-selected breaths are significantly different in many cases from expert-selected breaths, over 88% of the breaths selected by experts were also selected by BASSPRO. This suggests that because expert manual selection results in such a low sampling frequency of the data (∼10% of the data selected by BASSPRO), the variation between experts, and variation between experts and BASSPRO is actually a result of the method of manual selection missing data leading to undersampling. On the whole experts agree with BASSPRO as to what is quality breathing, but not all of the quality breaths are part of the expert selections on the individual level.

Together this supports the hypothesis that Breathe Easy demonstrates exquisite accuracy not solely because of the large number of breaths analyzed, but rather that BASSPRO’s filtering criteria extracts high quality breaths that includes the variety of calm, quiet breathing. Such representative variation present during calm, quiet breathing is exceedingly difficult to capture with manual selection.

Overall, the biological application of this software to our longitudinal Alzheimer’s study failed to reveal a consistent change over time, which suggests that forebrain APP expression alone is not a likely significant driver of respiratory dysfunction. This finding is somewhat surprising as forebrain streptozotocin AD models can drive changes in breathing rate^2, 3^. Further, it is well appreciated that forebrain injury from trauma or stroke can have lasting effects on breathing^28, 29^ and that forebrain seizures are capable of perturbing breathing^30^.

Of the results that were significant, we suspect that contributions of phenotypes previously reported in the control mice for this animal model to play a significant role^21, 31^. Given the variability of the control groups and the lack of any biologically significant development at 7 months off doxycycline, the differences reported at 7 months are most likely outliers. On the whole, there does not seem to be consistent evidence to support a direct link between the forebrain pathology of this model and pathological respiratory phenotypes.

Many additional features and outputs are being developed for future versions of Breathe Easy. The version presented here is a complete pipeline with calculation and analysis for the most frequently reported basic and refined respiratory variables. Future versions will aim to provide additional flexibility with user-defined breathing to include coughing and vocalization. We have also derived iterations of Breathe Easy for exploratory and data mining functions to investigate the dynamics of the waveform itself for predictive and diagnostic features. A component of these dynamics will include analysis of first and second derivatives of each component of the waveform for analysis of active vs. passive breathing and early/ late inspiration and expiration.

Using principal component analysis (PCA) of individual breath signals, it will be possible to interrogate traces for unknown features and phenomena that inform respiratory outcomes. Many groups are currently investigating the application of this analysis to raw waveforms, but the input to their analyses is limited to all breaths and noise from a recording or hand-picked sections of data^25, 32^. Additionally, several exciting avenues of breathing research are opening up that would benefit from modules to extract and merge other physiological signals such as ECG, EEG, EMG, and video for automated generation of ethograms describing an animal’s state as Breathe Easy does not currently segregate breaths by state.

## Methods

### Animal ethics statement

All animal studies were conducted at the AALAC accredited animal facilities at Baylor College of Medicine and all procedures were reviewed and approved by Baylor College of Medicine Institutional Animal Care and Use Committee under protocol AN-6171.

### Animal protocols

#### Breeding, genetic background, and maintenance of mice

Mice were maintained by mating a double Tg APP/tTA male on C57BL/6J background to a WT FVB/NJ female resulting in offspring that were FVBB6 F1. All four groups of mice used in this study were derived from this crossing: 1) wildtype (WT) FVBB6 (control), 2) Tg102_tetO- APPswe/ind (tetO) (control, MMRRC #34845^20^), 3) TgCaMKII*α*-tTA_LineB (CaMK) (control, Jackson Laboratories #3010^33^), and 4) Tg102_tetO-APPswe/ind; TgCaMKII*α*-tTA_LineB (Exp) (experimental). WT control mice are included as background genetic controls. Groups 2 and 3 represent single transgenic controls wherein transgenic insertions included in the experimental animals are present, but because only one of the transgenic alleles is included the resultant APP phenotype is not present. Group 4 is the experimental group where both transgenic alleles required for APP expression are present. Briefly, in the absence of doxycycline, tetracycline transactivator expressed under the CaMKII*α* promoter allows for the expression of the tetracycline-responsive chimeric mouse APP. This version of APP contains a humanized A*β* domain encoding the Swedish and Indiana mutations. Animals were housed on a 12h/12h light cycle with food and water ad libitum. Offspring were started on a Dox diet during the first postnatal week until 6 weeks of age when they were placed on regular chow. Animals were measured at multiple timepoints from ∼5 months after Dox withdrawal to ∼19 months after Dox withdrawal. There was no biologically significant difference in the weight or age of the mice in all groups at each recorded timepoint (Supplemental Tables 5C-D).

#### Experimental design

A total of 4 genetic groups were used in this study as outlined above. See Supplemental Tables 5A-B for the number of mice and Supplemental Table 5D for the average age of mice included for analysis for each timepoint, air exposure type, genotype, and sex. Briefly, we started with 11 WT controls, 12 tetO controls, 12 CaMK controls, and 11 Exp mice with sex evenly distributed for each group. The number of animals recorded does not always reflect the number of animals included in analysis as certain portions of the recording or entire recordings may not have been of sufficient quality. These cases lead to the mismatching in number of mice at those timepoints and conditions. Although halves of recordings may be included in analysis, our pipeline evaluates the results based on all breaths rather than averages, so all accepted breaths for each genotype at each time point for a given exposure are combined together to increase the power and accuracy of our analyses.

Each timepoint consisted of recording the mice for baseline room air breathing followed by either hypercapnic or hypoxic exposure. After a few days of recovery and reset, the same mice were recorded a second time for baseline room air breathing followed by the opposite challenge exposure, hypoxia or hypercapnia (Supplemental Fig. 2).

#### Whole body plethysmography

Whole body plethysmography on conscious, unrestrained mice was performed on ∼7 month to ∼21 month old adult mice^34^. Briefly, mice were habituated for 5 days and tested within 1 week of the last day of habituation. Habituation consisted of ∼5 minutes of handling, rectal temperature and body weight measurement, and 30 minutes in a plethysmography chamber^35^. On the day of testing, body weight and rectal temperature were recorded prior to the WBP experiment.

Animals were then placed in a water-jacketed, temperature-controlled (30°C), flow-through plethysmography chamber and allowed to acclimate for 30 minutes in hydrated (via an inline bubble chamber) room air gas (21% O_2_/ 79% N_2_). Following acclimation, the animal is recorded in room air for up to 1 hour for baseline measurements. After room air, the gas is switched to hydrated hypercapnic (5% CO_2_/ 21% O_2_/ 74% N_2_) or hydrated hypoxic (10% O_2_/ 90% N_2_) gas for a 20-30-minute exposure. Gas is then switched back to room air for up to one hour. Lengths of recordings are adjusted as needed based on animal activity to ensure sufficient collection of calm quiet breathing. The animal is removed from the chamber at the end of the last room air period and rectal temperature is recorded upon removal, and 30 minutes after removal. Body temperature from the beginning and end of the recordings are used to linearize body temperature over the duration of the recording in BASSPRO for use in Drorbaugh and Fenn- derived Bartlett-Tenney calculations^36, 37^. Exiting gases are dried via a nafion tube embedded in silicone desiccant prior to entry in gas concentration sensors.

#### Data collection

A Validyne DP45 differential pressure transducer was used to measure pressure changes between the animal chamber and a reference chamber. This signal was converted by a CD15 carrier demodulator. Oxygen and carbon dioxide concentrations were measured using AEI oxygen and carbon dioxide sensors and analyzers. Chamber temperature was measured using a ThermoWorks MicroThermo 2 thermocouple. All data was recorded via an ADI PowerLab analogue to digital recorder using LabChartPro in real time.

The raw signals produced during whole body plethysmography experiments are a time series of voltages corresponding to the sensor readings over the course of the recording session. In the experimental paradigm that we use, these sensors correspond to 1) the differential pressure produced within the recording chamber due to inhalation and exhalation via the countercurrent heating and cooling of respired air with a small contribution from compression and rarefaction outside and within the animal that is equilibrated as a function of airway resistance, 2) the temperature of the recording chamber, 3) the oxygen concentration of the gas leaving the chamber, and 4) the carbon dioxide concentration of the gas leaving the chamber. While the sampling frequency may vary, 1,000 Hz was used in the current data and has shown to be high enough frequency to permit high resolution and low risk of aliasing. The breathing waveform can be used to derive temporal and volumetric parameters related to breathing such as breath cycle duration, breathing rate, and an uncorrected tidal volume. The use of a temperature-clamped chamber to compare to body temperature provides a more accurate estimate of true tidal volume^36^. The remaining channels of data corresponding to oxygen and carbon dioxide concentrations can be used to calculate metabolic parameters such as oxygen consumption (*V̇_O_2__*), which can be combined with breathing parameters like ventilation (*V̇_E_*) to ascertain if changes in breathing are consistent with metabolic demand.

#### Consideration of data

The whole-body chamber, where the animal is housed during plethysmography experiments, is set up for near-balanced gas flow-through. During the experiment, the pressure fluctuations, largely due to heating and cooling of inspired and expired air respectively with a small contribution from compression and rarefaction of space inside and outside of the body in this chamber are compared to that of a reference chamber. The reference chamber is a duplicate of the containment device and does not house an animal during experimentation. Thus, any pressure differentials recorded between the experimental chamber and the reference chamber are from two sources. First, there may be slow changes in relative pressures due to inevitable imbalances with inflow and outflow. To guard against chamber over pressurization, as we run at a slightly higher inflow pressure, we connect the two chambers with a small gauge cannula to act as a low frequency band pass filter that prevents rapid breathing and movement pressure changes to transit the cannula. We also engineer a small leak to the atmosphere in the reference chamber to maintain the overall system at room atmospheric pressure. The final sources of pressure changes are respiration, respiration related behaviors (sniffing), and animal movement. This simple design allows for precise measurement of animal respiratory patterns. Optimization of flow characteristics ensures a stable baseline pressure waveform and minimizes noise in the data to facilitate software analysis. When this system is combined with oxygen and carbon dioxide sensors in line with the experimental chamber outflow, data can then be used to determine even more key variables including, but not limited to, oxygen consumption and carbon dioxide production, which enables the normalization of respiratory output to one or more metabolic parameters. Data resulting from an experiment using this or similarly designed custom systems are recorded using multi-channel voltage analog to digital recording devices such as PowerLab or LabJack (open source). We record at 1,000 Hz for highest resolution and minimal risk of aliasing, for each measured variable. Our software uses these multichannel voltage outputs as the starting point of processing. A robust and well-thought-out respiratory and metabolic system ensures high quality data with minimal artifacts for straight forward analysis of outcomes.

### Optimal plethysmography guidelines

While BASSPRO-STAGG can handle many common errors and mishaps that can occur during a plethysmography experiment to minimize data loss, there are general guidelines that, if followed, provide optimal recordings for analysis in BASSPRO-STAGG. Users need not re-perform experiments to abide by these guidelines. However, if protocols are updated to adapt to these considerations, then outcomes will most likely be far more accurate and reliable, and fewer experiments resulting in subpar recordings.

#### Adjustment of exposure protocol as needed

It is common practice for labs to develop a standard operating procedure for gas exposures in WBP. For example, 30 minutes of habituation, up to one hour of room air, 30 minutes of 5% CO_2_, and up to one hour of room air for challenge recovery, particularly for hypoxic challenges and the examination of the post-hypoxic response. While this standardization provides an ideal platform for ensuring consistency between recordings, unfortunately mice are not consistent. It is imperative that the operator performing experiments peruse the recorded sections of each condition of the experiment for quality breathing before proceeding to the next condition or section of the experiment. Some animals may take longer to reach a calm state, evident when movement artifacts are absent.

#### Habituation

While acclimation periods are most often included in a plethysmography run, habituation is less frequently included in experimental design. Rodents do not fully habituate to new environments within 30 minutes, particularly ones with significant differences from their home cages, e.g., size, foreign objects, and pressure. This lack of habituation results in increased noise as the animal explores its new housing as well as aberrant recording values related to the animal chewing on sensors and plugging gas intakes. To mitigate any contributions to noise derived from lack of habituation, we recommend habituating rodents once a day for at least 5 days where the last day of habituation is within a week of the plethysmography recordings^35^.

Habituation should include anything performed during the actual experiment, e.g., temperature recording, injections, handling, and housing inside a plethysmograph chamber.

#### Gas concentration calibration curve

To calculate gas concentrations accurately, the voltages recorded during the calibration phases of the experiment are used to assign the voltage to a known concentration for each recording session. This allows downstream calculations to be more accurate within the recording and between recordings. Often, however, only room air and the challenge gas for a given experiment are recorded during these calibration sessions. This creates a unique situation where a completed calibration curve with two values is only possible for one type of gas output. For example, if the user runs room air and 5% CO_2_ balance room air calibration sessions for a hypercapnic exposure, then only the CO_2_ calibration curve will have two points, one at 0% (room air) and one at 5% (5% CO_2_ balance room air). Furthermore, the accuracy of the gases used in these challenges is not 100%. Two different gas mixtures with an expected oxygen concentration of 21% may have slightly different readings. To mitigate these issues, we recommend that users 1) record calibration sessions for two types of challenge gases in addition to room air and 2) do not change the “zero” set point following the first gas. This allows the software to gather two calibration curve points for both types of gas that are analyzed, O_2_ and CO_2_. It also prevents user-derived voltage changes that deviate from the expected curves and would influence concentration calculations later in the recording.

### Breathing Analysis Selection and Segmentation for Plethysmography and Respiratory Observations – BASSPRO

The overall purpose of the waveform analysis program, called BASSPRO, is to import signal data and identify breaths within raw plethysmography data; calculate relevant respiratory parameters from those breaths; filter and group breaths into selections; combine metadata with the calculated respiratory parameters; and pass those data on to the statistics and graphing component of the program. This is accomplished using a variety of packages and series of operations that utilize the Python 3 programming language (packages summarized in Supplemental Table 6A).

#### Data import

Metadata are provided through an externally generated .csv file. Metadata represent three main categories: 1) periodic sampling during recording, 2) persistent animal metadata, and 3) initializing experimental data (Supplemental Figure 3). Periodic sampling during the recording are values obtained by the researcher during the recording that are specific to a given animal. These data are typically intrasession drug administration or body temperature where continuous body temperature is not always gathered in plethysmography experiments. Persistent animal metadata are all the details about the animal being recorded that do not change (e.g., the genotype, age, date of birth, sex, group (control or experimental), etc.). Initializing experimental data refer to information about the apparatus and conditions on the day of the experiment that may play a role in the outcomes measured. Examples of initializing data include the identification number for the plethysmograph (if more than one is available in the lab), the gas tank (if tanks are changed out while completing all recordings for a given experiment), body weight, room temperature, and room barometric pressure. If an animal is run more than one time, for example in a longitudinal study, then there will be initializing data for each run, but the persistent animal metadata will remain the same for every run of that animal. An example .csv with all required fields for BASSPRO can be accessed in our User Manual website hosted on our project GitHub page. The inclusion of as many factors as possible in the metadata, such as experimenter, specific gas tanks, and rigs, allows for the software-based evaluation of potential recording artifacts that may impact outcomes, e.g., a particular tank or WBP rig having significant error in the recorded data.

#### Breath selection and filtering

BASSPRO works by identifying every breath in a .txt file export of a WBP experiment followed by filtering of those breaths for desired qualities. The inputs to BASSPRO are all .txt files for an experiment and two or three configuration files, the number depends on the type of run the user chooses.

BASSPRO uses the two or three configuration files generated within the GUI to select and filter breaths including: 1) basic breathing parameters, 2) manual selections of breaths and 3) automated selection criteria. Basic breathing parameters are requirements that are used by BASSPRO to extract all breaths from the recording. The basic breathing parameters are settings that denote whether a peak found within the breathing waveform is a breath based on its characteristics, such as slope and duration, as well as using surrounding breaths to assess the presence of artifacts. A series of quality control steps are also implemented for the non-breathing signals such as chamber temperature and gas concentrations to identify data collection artifacts. If artifacts are present, then automated corrections may be applied and indicated in the log created by the run.

There are three ways BASSPRO-STAGG can be utilized to filter the identified breaths; manual selection, automated selection and filtering, and manual selection and automated filtering. The manual selection and automated selection criteria files inform BASSPRO on the three ways the user can run our program. A manual selection file includes breaths that are selected by the user from manual interrogation of the recorded plethysmography experiments. In this case, the user will perform most of the work of BASSPRO themselves and the Breathe Easy run will provide graphs and statistics. For automated selection, an automated selection criteria (autocriteria) file will be provided where the user determines what qualifications they deem appropriate for a breath to be considered calm, quiet breathing and which experimental conditions they would like to include in analysis. For example, the user could want only breaths if they are within a continuous group of at least 100 breaths that meet standards for calm, quiet breathing (based on the parameters of current and neighboring breaths) during the experimental conditions of room air and 5% CO_2_. To accurately calculate tidal volumes, the criteria for selecting the calibrated volume injections from room air and 5% CO_2_ may also be denoted in the autocriteria file. Default settings for a number of experimental paradigms come loaded as defaults in the automated settings subGUI for the user to select from. For manual and automated runs, the user provides their selections, and breathing based on both manual and automated filtering will be identified and included in the output. Calibration information from either manual or automated settings can be used, but if settings are provided in both configuration files, only the manual settings will be applied.

For any type of run, calibration periods must be identified for any sections where respiratory variables derived from Drorbaugh and Fenn, and Bartlett and Tenney formulae^36, 37^, which we call refined variables, are desired as an outcome. If calibration injections are not provided, then only basic breathing variables (breath cycle duration, uncorrected tidal volume, ventilatory frequency) can be calculated. An example of both a manual selection file and an autocriteria file are provided in the User Manual website found in the GitHub repository for this project. If variables from the experimental or animal information metadata are not input by the user, then most have default alternatives that are implemented so meaningful data can still be produced. Generation of these settings files and choices that can be made are more explicitly outlined in the User Manual website.

#### Export to STAGG

Multiple BASSPRO outputs are available including the main .json output that is utilized by the STAGG portion of this software suite, a .csv output enumerating all detected breaths that can be used to audit breath identification performance, and a .csv output providing aggregate summary values. The .json files are tables where each row is one breath that includes basic breathing variables, calculated respiratory variables, persistent animal metadata, initializing experimental data, and periodically sampled data during the recording. Each column is a value from one of those categories of data. One .json file is generated per animal per recording. A log file is also produced that can be used to verify settings used during a run, observe if irregularities were present in the signal files, or determine if corrections were applied during the processing of the files. Once all files have been created, they are stored in the output folder specified in the GUI in automatically generated folders that include the date and time of the run. Only the .json files are passed along to STAGG.

### Subprocesses of BASSPRO

BASSPRO performs all the filtering and selection of breaths as outlined above. However, there are a number of subprocesses that BASSPRO also performs during a typical run. Some of these processes are automatically performed and others are optional. It will be important for a user to familiarize themselves with the implications of each of these subprocesses as most are implemented to navigate a variety of common anomalies in plethysmography recordings. Also outlined is a more detailed explanation of how quality breathing is extracted from the raw waveforms.

#### Body temperature linearization

BASSPRO features a body temperature interpolation capability where body temperature measurements taken at the beginning, middle, and end of a session can be used to interpolate temperature values throughout the duration of the session for later use in calculating refined respiratory variables derived from Drorbaugh and Fenn, and Bartlett and Tenney formulae^36, 37^. Essentially, this function takes as many points as are available for temperature and creates a linear interpolator that connects those measurements throughout the recording. This artificially provides body temperature measurements for the entire recording rather than two to three distinct temperatures copied throughout the recording as is currently applied^6, 34^. Currently, this interpolation cannot be turned off and is applied in all cases where temperature is not continuously recorded.

#### Signal spiking adjustments for temperature, flow, and gas concentration channels

It is not uncommon for abrupt, high magnitude spikes or short square wave pulses to sporadically contaminate the voltage signals on some of the recorded channels. BASSPRO can apply a correction to the signal when this error is detected. For chamber temperature, when a value outside of the range of 18-35°C is detected for an otherwise good breath, then the chamber temperature is replaced either 1) with an interpolated value from surrounding temperature values that fall within the correct range, or 2) with a default value, which is set to 30°C if no temperatures from surrounding breaths fall within the desired range. Both the range of acceptable temperatures and the default value used for replacement can be changed in the basic settings subGUI in the *Rig Specifications* tab.

It is also possible that the chamber temperature can exceed the recorded body temperature of the animal, which will lead to errors in values calculated using the equations from Drorbaugh and Fenn, and Bartlett and Tenney^36, 37^. This particular error can be a result of drifting chamber temperature voltages, or aberrant body temperature measurements. In either case, the user has the option to exclude any breaths where the chamber temperature for that breath exceeds the body temperature for that breath. These breaths are removed by default but can be changed by editing the ‘High chamber temperatures inclusion’ variable in the *Additional Settings* tab of the automated settings subGUI.

For aberrant spiking in flow and gas concentration channels, the user can apply a smoothing filter, similar to flow smoothing filters commonly applied to WBP recordings. To apply the smoothing filter to flow and gas concentration channels, simply change the value for ‘Smoothing Filter’, which by default is 0=no smoothing on flow and 1=smoothing on flow, to “fg” in the basic settings subGUI in the *Breath Inclusion* tab. This setting by default only applies to the flow channel.

#### Automatic identification, selection of breaths, and basic respiratory variables

Breaths are identified through transitions between inspiratory and expiratory flows that are above their respective and tunable thresholds. Flow signal used for this identification is calculated as a simple first derivative of the original respiratory waveform as collected by a differential pressure transducer used for whole-body plethysmography recordings. When a flow is identified as crossing the threshold such that breathing has transitioned between inspiration and expiration (or vice versa), then the point at which the waveform crosses zero immediately preceding the threshold crossing is used as the time when the transition occurred. This permits identification of timepoints defining inhalation and exhalation for each breath cycle.

An optional filter can be applied (or implemented in later steps for automated selection) that utilizes the breathing frequency of neighboring breaths (Equation 1) and disagreement in amplitude of inhaled and exhaled volumes (Equation 2) to select breaths.

**Equation 1.** Percentage determination of neighboring breaths above a set breathing frequency where n is the instantaneous breath, w is ½ of the window surrounding the breath, F is the instantaneous breathing frequency, and X is the threshold set by the user (recommended as 500 for mouse breathing).

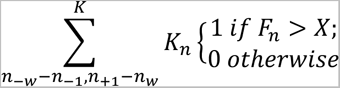

**Equation 2.** Calculation of inhalation/ exhalation volume disagreement.

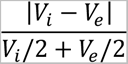

During the initial identification of breaths, several parameters are calculated for each breath, which are available as part of the output. Some have been identified as useful for automated selection criteria as well. The candidate breaths that are initially identified are then available for manual or automated selection. Manual selection will place all breaths contained within manually-defined intervals within the output. Automated selection relies on a series of filters to refine the list of candidate breaths into the set of breaths contained within the largest continuous interval for each experimental condition as well as the subset of those breaths that meet criteria for “calm, quiet breathing”.

Parameters that are defined for each breath include the duration of inhalation and exhalation, the total breath cycle duration, the instantaneous breathing rate, uncorrected inhaled and exhaled tidal volumes, peak inspiratory and expiratory flows, and scoring for featured breathing such as being an apnea or sigh (1=True, 0=False). Apneas can be defined by duration relative to neighboring breaths (at least twice as long recommended for mice) and minimum length of apnea (0.5 seconds recommended for mice), and sighs are defined as breaths with increased tidal volume relative to neighboring breaths (at least twice the amplitude is recommended for mice). These definitions can be changed based on user requirements, but these defaults represent standards of the field.

Irregularity scores are also calculated for these basic respiratory parameters relative to the preceding breath. The calculation for irregularity scores is shown in Equation 3 where X is the value for which an irregularity score is being calculated and N is the number of the current breath for which the irregularity score is being calculated.

**Equation 3.** Irregularity score calculation.

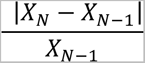

To facilitate identification of calm quiet breathing, additional parameters are also calculated as Boolean True/ False values based on the underlying parameters value relative to the thresholds set in the autocriteria file.

#### Automated sectioning by comments

The user has the option of using comments inserted during the plethysmography recording as experimental landmarks that BASSPRO can use to divide the recording. The initial release of BASSPRO is designed to work with signal files exported from LabChart and can parse timestamps placed using the LabChart software. Any recording that is exported with comments in the style of the LabChart text file should also work given that comments in the autocriteria file match those found in the recording. To use comments that are already contained in the signal files, the user will include those as text strings in the autocriteria file, which is only required if the user desires automated selection of breaths from their signal files. Each comment within the autocriteria file is used to define the beginning of an experimental condition whereas the ending of an experimental condition is defined by the next comment in the LabChart file that is also listed in the autocriteria file, or by the end of the LabChart file if no suitable comment is found.

The first division of experimental conditions is into “verified conditions” that meet the basic inclusion criteria based on their timestamps and gas concentrations. The user may specify offsets from beginning or ending of an experimental condition as well as minimum and maximum O_2_ and CO_2_ concentrations that can be used to refine the appropriate window for selection of breaths (i.e., to account for and exclude ramping gas concentrations during transition periods). These parameters are intended to be set by the user to define objective criteria for inclusion. Verified experimental conditions that pass the inclusion criteria for the condition are passed on to the next set of filters, verified breathing.

#### Refined variable calculation

Several respiratory parameters are automatically extracted from each identified breath whether the breath is assigned a corresponding calibration period or not. These include, but are not limited to, breath height, breath width, breath frequency, breath cycle duration, and uncorrected tidal volume. Drorbaugh and Fenn-derived Bartlett-Tenney corrections are used to calculate more refined estimates for respiratory volumes and metabolism^36, 37^. To make those calculations, calibration injections must be included so accurate volumes of gases inhaled and exhaled, and baseline concentration voltages for oxygen and carbon dioxide concentrations can be calculated. Variables that are calculated in this nature include corrected tidal volume (V_T_), minute ventilation (*V̇_E_*), oxygen consumption (*V̇_O_2__*), and ventilatory equivalents for oxygen (*V̇_E_*/*V̇_O_2__*).

It should be noted that the nature of calculating these values introduces a division transform problem. This problem impacts the aggregate value calculated for an animal for a gas type or condition. For example, you can either calculate these variables using the average of recorded values for the animal during the gas exposure, or you can calculate these variables for each breath and average at the end. These methods will yield similar, yet slightly different results with their divergence typically increased as breathwise variation in breathing variable values increases. With STAGG, our statistical model is built on the analysis of individual breaths, with the ability to apply rank preserving transformations to the data to address non-normality in the statistical model. We utilize the individual observations for the statistical analysis performed in STAGG in order to account for the variability in individual measurements present in the observed data. Using the individual breaths for analysis also increases the statistical power of hypothesis tests relative to using aggregated mouse values. However, some users may prefer the traditional approach of derivation of refined variables from pre-aggregated components, and BASSPRO can provide an optional output file that makes these available.

Equations 4-9 elaborate the calculations made for each breath during a standard BASSPRO run.

**Equation 4.** Vapor pressure calculation where T is the temperature (Kelvin) of the air where vapor pressure is desired.

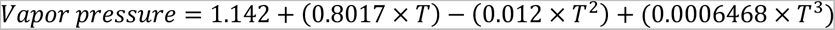

**Equation 5.** Corrected tidal volume (V_T_) calculation with Drorbaugh and Fenn-derived Bartlett-Tenney correction where A is voltage from the breath, B is average voltage of calibration injections, C is nominal volume of calibration injections, D is body temperature (C), E is room barometric pressure (mmHg), F is vapor pressure of the chamber, G is chamber temperature (C), and H is vapor pressure inside the mouse.

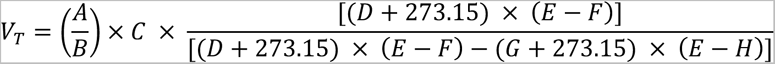

**Equation 6.** Minute ventilation calculation using corrected tidal volume (Equation 2).

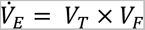

**Equation 7.** Oxygen consumption (*V̇_O_2__*) calculation where oxygen in is taken from voltage during calibration, and flowrate is recorded as initializing experimental data in liters per minute (LPM) and converted to milliliters per minute (mLPM) in the equation. A is flow rate (LPM), B is oxygen concentration in (%), and C is oxygen concentration out (%).

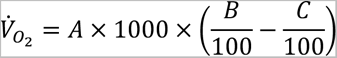

**Equation 8.** Carbon dioxide production (*V̇_CO_2__*) calculation where carbon dioxide in is taken from voltage during calibration, and flowrate is recorded as initializing experimental data in LPM and converted to mLPM in the equation. A is flow rate (LPM), B is carbon dioxide concentration out (%), and C is carbon dioxide concentration in (%).

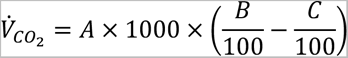

**Equation 9.** Ventilatory equivalents for oxygen (*V̇_E_*/*V̇_O_2__*) calculation using minute ventilation (Equation 3) and oxygen consumption (Equation 4).

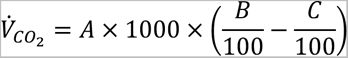

### STAtistics and Graph Generator – STAGG

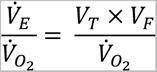

The overall purpose of the STAGG platform of Breathe Easy is to statistically analyze and graph all of the desired variables for a given experiment. The flexibility coded into this component allow the user to analyze almost any desired respiratory variable along with any metadata external to the WBP recordings, like weight, age, or sex. Additional quantified data from paired behavioral assessments or even immunohistological cell counts can also be included. STAGG is comprised of a series of R scripts that process the data from BASSPRO. All packages can be found in Supplemental Table 6B.

#### Data import and statistical analysis

First, the .json output of BASSPRO as well as the STAGG configuration files are loaded. Using the rjson package the data import script reads in each .json file one at a time and prints to the user the filename currently being imported as a progress bar. Once a .json is imported, it is added to the end of the previously imported .json to create a large tibble containing all data from every file. STAGG requires 3 configuration files: 1) variable configuration, 2) graph configuration, and 3) other configuration. The variable configuration file outlines all the statistical settings for a given experiment where independent, dependent, and covariate variables are assigned. This file will also house permutations or settings specific to the variables that are set by the user in the STAGG settings subGUI, which includes things like transformations, minimums and maximums of the y-axis, and the generation of Poincaré plots. This file will also include ‘Alias’ values that may be assigned by the user. For example, instead of VO2pg appearing on a graph, the user may instead prefer Oxygen Consumption. In this case, the user can reassign the name for VO2pg by replacing VO2pg in the ‘Alias’ column with Oxygen Consumption. The graph configuration file contains instructions on how to plot the dependent variables selected on the graph. For example, if you want to see your dependent variables graphed by air condition and sex, then you would select air condition as xvar and sex as pointdodge. Finally, the other configuration file outlines optional graphs that are selected by the user. In optional graphs you can graph variables that may not be respiratory variables, but they are metadata that you want to plot for the experiment. You can also plot your respiratory variables with different settings than those in the primary model, which are selected in the graph settings. Settings for optional graphs follow the same guidelines as those for variable and graph configuration. Explicit instructions and more details for assigning values and making selections for these files are included in our User Manual website hosted on our GitHub page for this project.

Assignments for independent, dependent, and covariate variables are employed to fit separate linear mixed effects models from the data for each of the designated dependent variables using the *lmer* function from the *lme4* package in R as well as any user-selected transformations for these variables. The independent variables are included in the model as fixed effects, along with an interaction variable which involves all designated independent variables. On the other hand, only the main effects for the covariates are included in the model as fixed effects. The unique mouse identification number (MUID) is included in the model as a random effect. Once these are set, a loop processes each of the dependent variables one at a time with the assigned independent and covariate factors. Results of the LMEM then undergo post hoc analysis with a Tukey’s honestly significant difference (HSD) test using the *glht* function from the *multcomp* package, in which pairwise differences between different interaction groups are tested for statistical significance. The outputs of this final test are compiled into a single table. This is then filtered to keep only biologically relevant significant results, i.e., only results which are both statistically significant and for which there is only a single difference amongst the combined group levels of the interaction variables. These results are used during the graph generation process to place asterisks that signify significant differences where appropriate. Full tabular results from the mixed effects model and the Tukey post-hoc tests are saved in an Excel spreadsheet where each dependent variable is given a tab in the sheet that holds results for each comparison made, and QQ and residual diagnostic plots for each individual mixed effects model are saved in the same StatResults folder.

For optional graphs, the primary LMEM model are used for statistics unless the response variable chosen either 1) is not included in that model or 2) is not a continuous variable. If 1, then a new LMEM is run with the selected graphing options used in that model as independent variables. If 2, then an ANOVA is used to generate statistics for that graph. Results of these statistical tests are saved in a separate statistics folder from the primary LMEM.

#### Graph generation

All dependent variables assigned by the user are graphed with statistical analysis results added to each. Settings from the GUI that are stored in the graph configuration file designate how the variables are plotted, i.e., which variables the dependent variables will be graphed against.

Once these are assigned, scatterplots are generated via the *ggplot* function which show each dependent variable plotted against the selected independent variables or covariates at the individual mouse level. If there are significant results in the Tukey output, then asterisks are added to the appropriate group based on the location identified for each comparison and dotted lines are added to denote the groups that are statistically significantly different from one another. After each component has been added to the graph, the graphs are saved as either .svg or .jpeg files in the designated output folder.

#### Optional graphs

The final graph making script allows the user to choose any combination of independent and/ or dependent variables for plotting. This can include anything from body weight and body temperature, to plotting the dependent variables by different independent or dependent variables. This flexibility also opens the possibility for graphing results from behavioral studies and other tests performed outside of the plethysmography recording.

#### R Markdown and ReadMe

STAGG is completed by the run of an R markdown script that grabs all .svg or .jpeg files that are present in the selected output folder and compiles them into an .html page where the user can view all graphs generated in one place. This feature is ideal for the user because it allows perusal of data without needing to open multiple individual files.

In addition to an .html file with all figures produced, a ReadMe word document is produced that houses all of the settings the user selected for that particular run of STAGG. It eliminates all variables that were not assigned a role in the variable configuration file to make perusal easier for the user. This ReadMe file is ideal for keeping track of different runs of the software and taking notes on adjustments or changes that should be made from that particular run.

### Graphical user interface – GUI

The GUI is a PyQt5-built program that integrates BASSPRO and STAGG into a portable desktop application. The GUI’s design follows the standard workflow of plethysmography analyses. Lateral panels display the paths of selected input for both modules, providing immediate feedback to the user on their selections. At the bottom of the GUI is the hangar display, where more explicit feedback as well as terminal output is posted for user review. On either side of the hangar display, the user can launch either module of Breathe Easy, or launch them sequentially. The GUI addresses 3 main issues with running BASSPRO-STAGG: 1) portability, 2) processing power/ speed, and 3) settings and formatting input.

Various subGUIs are used to provide specific input required by BASSPRO or STAGG. They are designed with the user in mind in that they prompt the user to input settings that are coded within BASSPRO-STAGG without needing to open, understand, or edit the code or configuration files directly. These subGUIs are easy to navigate and walk the user through all definitions of parameters required as well as recommended settings based on our experience. There are five total subGUIs: 1) basic BASSPRO settings, 2) manual BASSPRO settings, 3) automated BASSPRO settings, 4) metadata settings, and 5) STAGG settings. Explicit instructions on using the GUI and subGUIs and expectations for their inputs can be found on the User Manual website hosted in our GitHub repository.

#### Portability

A critical feature of the BASSPRO-STAGG application is its portability. The program employs a variety of languages, libraries, and packages from the Python and R universes. To accommodate novice users that are not skilled in setting up Python and R environments, a distributable pre-compiled package was created permitting installation as easy as downloading an archive file, unzipping the contents, and running the executable file contained within. Nuitka and R-Portable were utilized to create packaging that allows the user to run the application without needing to download or run Python or R. To facilitate this portability, the GUI creates default directories and identifies relative paths that allow the program to run independent of local environments. Currently, our GUI only works with PC machines. However, the code can be run manually on a Mac, and we are in the process of creating a Mac executable to open and run the GUI.

#### Parallel processing

Another critical global feature of the GUI is its implementation of parallel processing. The GUI uses the *concurrent.futures* module to spawn separate processes for every signal file, allowing it to implement asynchronous execution for signal file interrogation. The user can choose the number of CPU cores they wish to devote to BASSPRO, which allows them to continue experiencing adequate performance while running other programs on the same computer. This parallel processing greatly reduces runtime, depending on configuration.

If parallel processing is not chosen by the user, then BASSPRO defaults to threading the signal files sequentially through BASSPRO. On average, the interrogation of a single signal file representing a 1.5 to 2 hour-long plethysmography run takes two to three minutes on contemporary computer hardware. Running 16 signal files through BASSPRO one at a time would take 32-48 minutes. Running those same 16 signal files through BASSPRO using parallel processing, with at least eight processing cores, can finish the task within six minutes. On average, we see an 81.8% reduction in run time with parallel processing compared to serial processing. Note that the majority of consumer computers are limited to eight processing cores. This is a dramatic contrast to the 50 minutes-3.15 hours spent by experts to manually select breaths from the same 16 files. Either option, parallel or sequential processing, is an improvement on the time and labor cost. It should be noted that although all cores can be sent instructions during the parallel processing procedure, with contemporary machines, there should be enough processing power to continue operating other applications while BASSPRO-STAGG is running. This said, depending on configuration, running other applications during BASSPRO- STAGG can potentially slow the processing time as the full processing power of each core would not be fully dedicated to the pipeline.

Currently, BASSPRO is the only module of the pipeline that utilizes this feature. Utilizing parallel processing for STAGG in its current form would not work or provide any benefit as there are many steps requiring sequential reading and writing throughout the STAGG module that make parallel processing impossible. However, once a dataset has been run through STAGG, one of the outputs is an R environment created just after all of the input .json files are read into R. If an R environment is selected as the start point for STAGG rather than all of the .json files, then there is an 82.2% reduction in run time. So, there is still an opportunity to increase the speed at which STAGG runs without parallel processing.

### Sub-GUIs

Each subGUI discussed below provides specific input required by BASSPRO or STAGG. They are designed with the user in mind in that they prompt the user in layman terms to input settings that are coded within BASSPRO-STAGG in a less intuitive fashion. These subGUIs are easy to navigate and walk the user through all definitions of parameters required as well as recommended settings based on our experience. There are 5 total subGUIs: 1) basic BASSPRO settings, 2) manual BASSPRO settings, 3) automated BASSPRO settings, 4) metadata settings, and 5) STAGG settings. Not all the subGUIs are required for each type of run (i.e., manual vs automated). Each of the subGUIs produces a uniquely formatted .csv file that is sent to the appropriate program to provide the required instructions. Please see our User Manual website for more information and screenshots.

#### Basic BASSPRO settings subGUI

There are four groups of data and user definitions input in the basic parameters subGUI. These groups are separated into four tabs at the top of the subGUI with a fifth tab that summarizes all of the selections made in the other four tabs. Please see the User Manual website for more information on choosing settings in this subGUI.

Breath calling settings describe minimum quantitative guidelines for a peak in a recording to be considered a breath. This does not define the breath as calm, quiet breathing, rather just identifies that peak as breathing. Breath calling also defines featured breathing that may not pass quality breathing filters set later, like apneas and sighs. Defining these features allow the user to quantify these kinds of breaths separately from calm, quiet breathing. Rig specifications are where settings specific to your plethysmograph will need to be detailed for use in respiratory calculations from the raw waveform. Breath inclusion filters are very basic filters that can be set for an initial pass of exclusion of breaths. Generally, the defaults set here are recommended for a first pass through data. Output settings allows the user to indicate in which formats, other than the .json files, they would like the output of BASSPRO saved. Options here can be helpful for perusing the raw data in Excel.

#### Manual BASSPRO settings subGUI

The manual settings subGUI streamlines the creation of the manual selection file needed for BASSPRO to analyze signal files with breaths that have been manually selected. The subGUI requires the user to select a folder containing the .txt files produced by exporting LabChart DataPad views. If a user is making selections with a software other than LabChart, then those selections need only be formatted to match the style and headings of DataPad selections found in LabChart for this subGUI to be used. Alternatively, they can directly generate a manual selections file using our example file provided within the documentation for the User Manual.

These exported files are collated into a table presented in the subGUI window, allowing the user to review their selection of signal files. The user specifies the type of experimental condition applied during the plethysmography runs represented by the chosen signal files. Based on the user’s selection of experimental condition, the subGUI provides default settings for BASSPRO to use during its analyses. These settings are presented as a table in the subGUI that the user can review and edit, as needed. Upon merging the collated DataPad views with the settings chosen, the subGUI creates the manual selection file by assigning the appropriate settings to each row of the collated DataPad views. This table is then exported to BASSPRO for analysis. There are a number of assumptions made based on the file names for the recording files and exported DataPads. Please see the User Manual website for more information on how to export, guidelines for naming signal and settings files, and what options are available for manual selection.

#### Automated BASSPRO settings subGUI

The automated settings subGUI facilitates the creation and modification of the automated criteria (autocriteria) file needed for BASSPRO to analyze signal files by identifying breaths and features, filtering those breaths, and calculating output measures. The subGUI only asks the user to specify the experimental paradigm relevant to the signal files they wish to analyze. Their selection will populate the subGUI with values based on the default settings for that experimental condition. The user can also choose to load a previously made autocriteria file that will populate the subGUI with the settings defined therein. Alternatively, the user can derive all new settings based on comments pulled directly from the signal files. This option requires more work and should be done with caution and help from our User Manual. Automated selection settings vary depending on the section of the plethysmography run, so each setting will have several values, one for each of the sections. The user can review the values of all sections for each setting and edit those values as needed. These modified settings can then be saved as a new autocriteria file that will dictate how BASSPRO detects and filters the breaths in the signal files. Once generated, these settings are collated into a .csv file that is exported to BASSPRO. Suggested default settings can be found in a table hosted in our User Manual website.

#### Metadata settings subGUI

The metadata settings subGUI was designed to streamline the codification and reorganization of variables found in the metadata. This subGUI allows the user to rename variables, convert continuous variables into categorical variables, and create novel groups of variables. It modifies the metadata intended as BASSPRO input without altering the original format, instead creating a new file that can be used for subsequent BASSPRO analyses. Continuous variables can be difficult to visualize in a graph, so creating bins for ranges of values can allow the user to convert continuous variables into categorical variables for an easier to interpret graphical output, e.g., one month and two months instead of P30, P32, P59, and P64. Novel groups of variables allow the user to group animals together into meaningful biological groups for easier comparison even if the values for their grouping are different, e.g., Control and Experimental instead of Tg, WT, Het. Each of these permutations of the data are added to the existing data, which allows the user to re-load the same data files and perform different runs with the same dataset based on the exact question they would like to answer.

#### STAGG settings subGUI

The STAGG settings subGUI was designed to facilitate the configuration of statistical and graphical parameters used in STAGG. It presents the user with variables sourced from the metadata and analysis parameters provided as BASSPRO input as well as variables derived from BASSPRO output. This collation allows the user to assign statistical and graphical roles to variables of interest by designating them as independent, dependent, or covariate for statistical analyses and choosing how they are presented in the figures produced by STAGG. This interface also permits renaming of variables, allowing the user to create new naming conventions for a variable that they would like to be displayed on a graph, e.g., Ventilatory Frequency instead of VF. Please see our online User Manual for more information and help understanding some of the statistics employed in our software.

### Machine requirements

General machine requirements to run BASSPRO-STAGG vary based on the size of the user’s dataset. In general, we recommend twice as much RAM as the size of the full dataset. The user should have a minimum of 5GB disk space to download and install and a minimum of 4 GB DDR3 RAM, which should be sufficient for a small dataset. We recommend 50 GB disk space to download, install, and store datasets with 16 GB DDR3 RAM. If datasets are highly complex, i.e., more than 20 combinations of independent factors and more than 50 recordings, then 32 GB+ DDR3 RAM are ideal. Outside of this recommendation, minimum and recommended operating systems and system architectures are described in Supplemental Table 7.

Later STAGG development was largely affected by the demands of working with very large datasets. The size of the data table during importation, the size of the vectors produced during modeling, and the number of pairwise comparisons each represented bottlenecks in performance. STAGG loads the BASSPRO output into a single table that is actively maintained in RAM while the program is building it during importation. A crude indication of whether or not the computer running STAGG has sufficient RAM to handle the dataset is to compare that RAM to the collective size of the input files. A computer with 16 GB of installed RAM cannot create a table collating data from input files that collectively take up more than 16 GB of memory. There are two workarounds: 1) use a computer with more RAM, or 2) reduce the size of the input files by removing data that will not be used in the analysis. The removal of data could be removing a continuously recorded channel or metadata that is not used.

Similarly, sufficient RAM is needed to maintain the vector(s) created when STAGG models the data. This issue is again related to the size of the dataset with which STAGG is working.

However, an additional factor at play is the number of comparisons made in the model. You may have insufficient RAM to maintain a few large vectors or many smaller vectors. Additionally, one of the libraries used to build the model caps the number of pairwise comparisons at 1,000. These related, but separate, issues were addressed by only modeling comparisons of interest, which are defined as differing at only one level of the independent variables being compared.

This restriction has proven to be an acceptable and encompassing definition for biologically relevant comparisons and, therefore, an acceptable filter.

### Data Availability Statement

All of our source code as well as a downloadable launcher for the GUI is available in our public GitHub repository (https://github.com/MolecularNeurobiology/Breathe_Easy). Within the ReadMe are multiple links to our online documentation (Sphinx and GitHub Pages). We highly suggest first time users to install and launch the GUI, then walk through the vignette provided that includes an example dataset and settings files.

Raw data for our complete Alzheimer’s dataset is too large to store on Zenodo, which is our default data hosting site. However, we will share the data via cloud-based services upon request. Please send correspondence to Russell Ray at for requesting access.

## Acknowledgements

We thank J. Sun, V. Martinez, and F. Saldana for technical support. Funding from NIH and NSF. The funders had no role in study design, data collection and analysis, decision to publish, or preparation of the manuscript.

## Funding

S. Lusk: NIH R01HL130249, 44617-S4 (Diversity Supplement) and NIH 1F32HL160073-01A1

G. Allen: NSF NeuroNex-1707400

J. Jankowsky: NIH RF1 AG054160

R. Ray: NIH R01 HL130249, NIH R01 HL161142, McNair Foundation

This project was supported by the Mouse Metabolism and Phenotyping Core at Baylor College of Medicine which is subsidized through Baylor College of Medicine’s Advanced Technology Cores and through funding from NIH (UM1HG006348, R01DK114356)

## Contributions

Experimental design (S.L., C.S.W., J.J., R.R.); software development (S.L., C.S.W., A.C., A.T.H., S.F.); animal preparation (J.J.); data collection (S.L., R.R.); analysis design and implementation (S.L., C.S.W., A.C., A.T.H., S.F., G.A., J.J., R.R.); manuscript preparation (S.L., C.S.W., R.R.).

## Ethics Declaration

The authors declare no competing interests.

## Supplemental Information

**Supplemental Figure 1.**
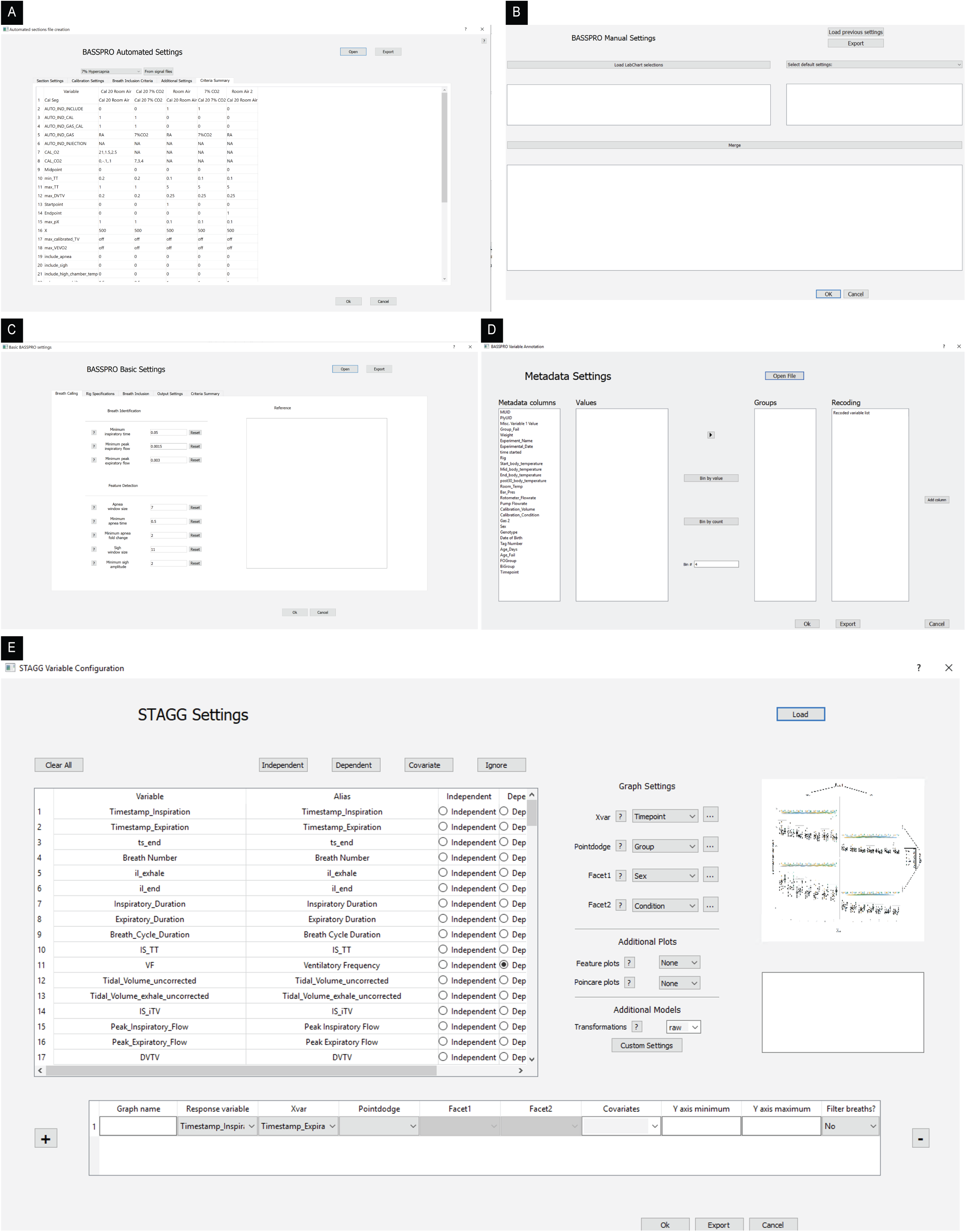
Screenshots of each of our subGUIs that can be launched from our main GUI: A) automated settings, B) manual settings, C) basic settings, D) metadata configuration, and E) STAGG settings.

**Supplemental Figure 2.**
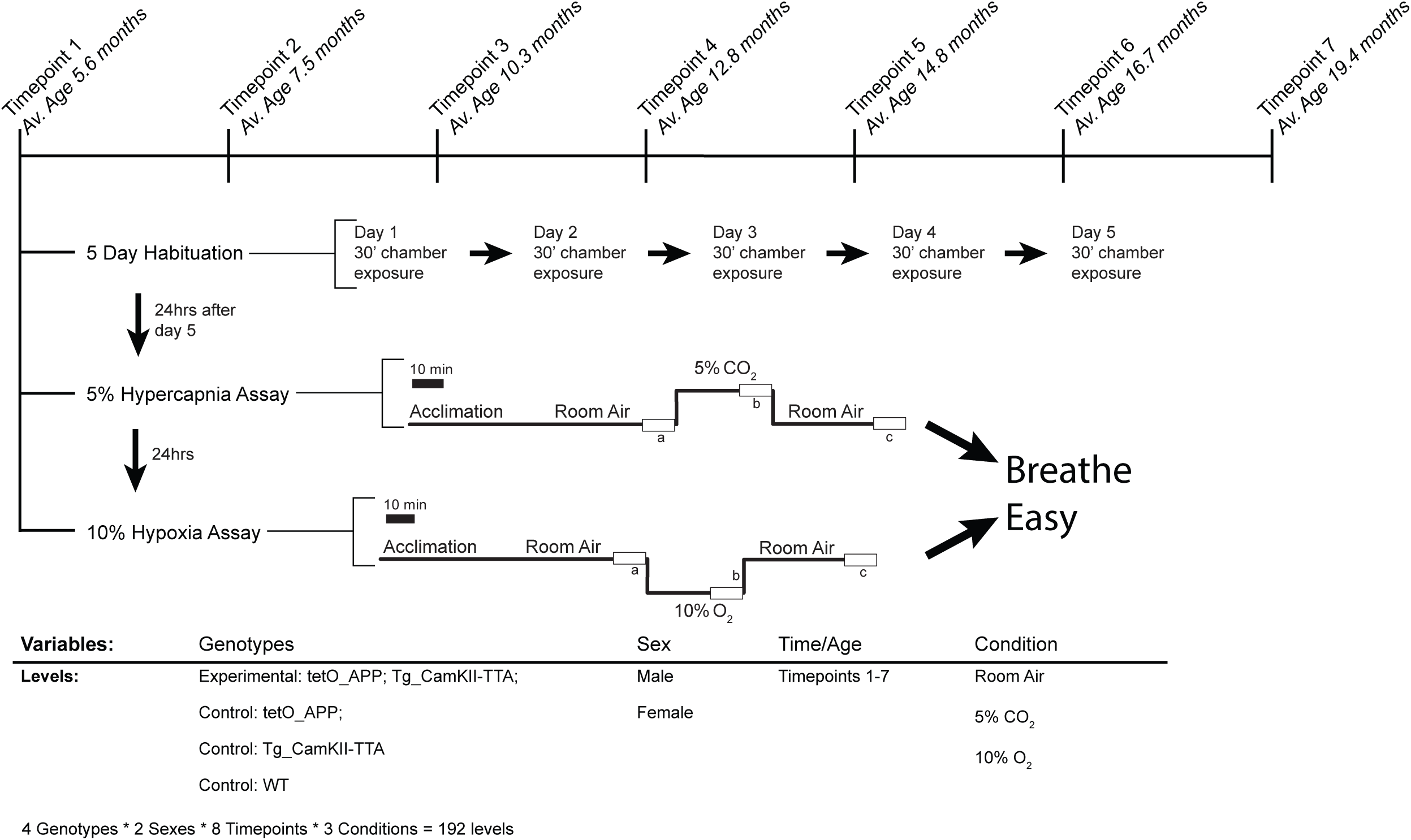
Schematic of our longitudinal Alzheimer’s study where average age at each timepoint across the length of the experiment is shown followed by a detailed protocol followed with each animal at each timepoint. Also provided is an explanation of variables included in analyses for this study.

**Supplemental Figure 3.**
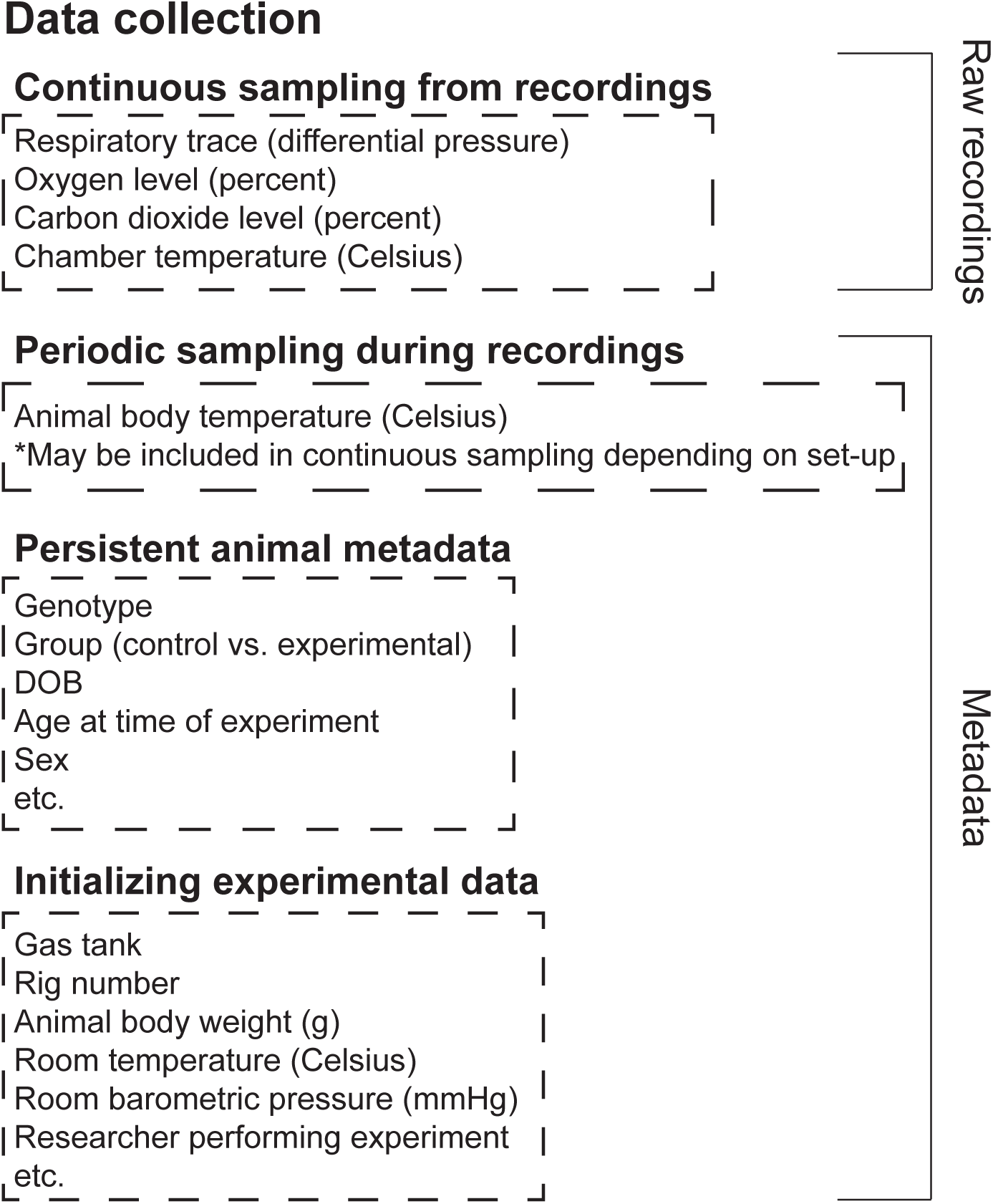
Schematic of variables collected for a plethysmography experiment that can/ must be included in Breathe Easy for analysis.

**Supplemental Table 1.** Non-exhaustive literature review of reported respiratory variables from 1998-2022 from a variety of laboratories across the world. Values were taken from graphs as most publications did not report raw values. Only values for control, adult mice were used to populate this table.

| Ref | Year | *Original units reported* |  |  | *Values reported after conversion* |  |  |
| --- | --- | --- | --- | --- | --- | --- | --- |
|  |  | Respiratory rate/<br>Ventilatory frequency | Tidal volume | Minute volume/<br>Ventilation | Respiratory rate/<br>Ventilatory frequency<br>(breaths min.<sup>-1</sup>) | Tidal volume<br>(mL breath<sup>-1</sup> g<sup>-1</sup>) | Minute volume/<br>Ventilation<br>(mL min.<sup>-1</sup> g<sup>-1</sup>) |
| 1 | 2022 | breaths min.<sup>-1</sup> | mL breath<sup>-1</sup> g<sup>-1</sup> | mL min.<sup>-1</sup> g<sup>-1</sup> | 55-275 | 0.002-0.014 | 0.01-3 |
| 2 | 1998 | Not explicitly reported. | μL g<sup>-1</sup> | μL sec.<sup>-1</sup> g<sup>-1</sup> | 73-162 | 0.0065-0.0085 | 0.9-1.02 |
| 3 | 2019 | breaths min.<sup>-1</sup> | mL breath<sup>-1</sup> g<sup>-1</sup> | mL min.<sup>-1</sup> g<sup>-1</sup> | 125-260 | 0.004-0.013 | 0.01-3 |
| 4 | 2002 | breaths sec.<sup>-1</sup> | μL g<sup>-1</sup> | mL min.<sup>-1</sup> g<sup>-1</sup> | 150-180 | 0.0057-0.0061 | 0.75-1.1 |
| 5 | 2021 | breaths min.<sup>-1</sup> | mL kg<sup>-1</sup> | mL min.<sup>-1</sup> kg<sup>-1</sup> | 150-600 | 0.008-0.013 | 1.75-5 |
| 6 | 2012 | breaths min.<sup>-1</sup> | mL | mL min.<sup>-1</sup> | 170 | 0.011* | 2.18* |
| 7 | 2017 | breaths min.<sup>-1</sup> | mL | mL min.<sup>-1</sup> | 175-190 | 0.0036-0.0054 | 0.54-0.71 |
| 8 | 2010 | cycles min.<sup>-1</sup> | μL g<sup>-1</sup> | mL min.<sup>-1</sup> g<sup>-1</sup> | 180 | 0.0064-0.007 | 1.3 |
| 9 | 2016 | breaths min.<sup>-1</sup> | NR | NR | 185-210 | NR | NR |
| 10 | 2015 | breaths min.<sup>-1</sup> | PO | NR | 187-211.9 | PO | NR |
| 11 | 2013 | breaths min.<sup>-1</sup> | μL g<sup>-1</sup> | mL g<sup>-1</sup> | 231-253 | 0.0049-0.0061 | 1.2-1.38 |
| 12 | 2020 | breaths min.<sup>-1</sup> | NR | NR | 250 | NR | NR |
| 13 | 2017 | breaths min.<sup>-1</sup> | mL breath<sup>-1</sup> g<sup>-1</sup> | mL min.<sup>-1</sup> g<sup>-1</sup> | 275-300 | 0.12-0.14 | 3.5-4.5 |
| 14 | 2018 | breaths min.<sup>-1</sup> | mL breath<sup>-1</sup> | mL min.<sup>-1</sup> | 400 | 0.012-0.015 | 5 |
| 15 | 2018 | breaths sec.<sup>-1</sup> | NR | NR | 420 | NR | NR |
| 16 | 2011 | NR | NR | mL min.<sup>-1</sup> g<sup>-1</sup> | NR | NR | 3 |
| 17 | 2013 | NR | NR | μL | NR | NR | 3.11 |
| 18 | 2006 | PO | PO | PO | PO | PO | PO |
| 19 | 2010 | PO | PO | mL min.<sup>-1</sup> | PO | PO | 1-1.6 |

NR = not reported and PO = percentage only. PO also includes percent change, arbitrary units, and delta values.
\*Where units reported do not match the standard of Breathe Easy, conversions were made using reported body weight values or standard conversions to match the units to those of Breathe Easy to allow for easier across-publication comparisons. If body weight was not reported, an \* marks those values and average body weight was assumed to be 27.5 g.

**Supplemental Table 2.** Feature comparison of commercially available products to Breathe Easy. This table was generated using publicly available information from each of the companies’ websites.

| **Manufacturer:**<br>**System/Software:** | **<u>Breathe Easy</u>** | **DSI**<br>***Finepointe*** | **DSI**<br>***Ponemah*** | **ADI**<br>***LabChart*** | **EMKA Scireq**<br>***iox2*** |
| --- | --- | --- | --- | --- | --- |
| **Features** |  |  |  |  |  |
| *Free/ Open source?* | Y | N | N | N | N |
| *Data collection* | Designed around ADI Powerlab data collection but can use outputs from other systems through conversion to .txt file input standard | Y (limited to Finepointe) | Y (limited to Ponemah) | Y (also has signal import if in compatible format) | Y (must purchase conversion software to use with recordings performed on other systems) |
| *Breath detection* | Y | Y | Y | Y | Y |
| *Calculation of basic breath parameters (volume, frequency, flow, etc.)* | Y | Y | - | Y (frequency and volume) | Y |
| *Detection of breathing features (apnea, sigh, cough)* | Y (apnea, sigh) | Y | - | - | - |
| *Calculation and normalization with metabolic parameters (VO<sub>2</sub>, VCO<sub>2</sub>)* | Y | Y | - | Y | - |
| *Manual sub-selection of data for analysis* | Y | Y | - | Y | Y |
| *Automated selection of data for analysis (basic per breath)* | Y | Y | - | - | Y |
| *Automated selection of data for analysis (filtered quality breathing from desired conditions only)* | Y | - | - | - | - |
| *Summary figures* | Y | Y | - | - | Y (requires separate software purchase) |
| *Summary statistics* | Y | Y | - | - | Y (requires separate software purchase) |
| *Statistical comparisons between groups* | Linear Mixed Effects Model with Tukey Post-hoc; ANOVA for some metadata comparisons when appropriate | ANOVA | - | - | - |

**Supplemental Table 3.**
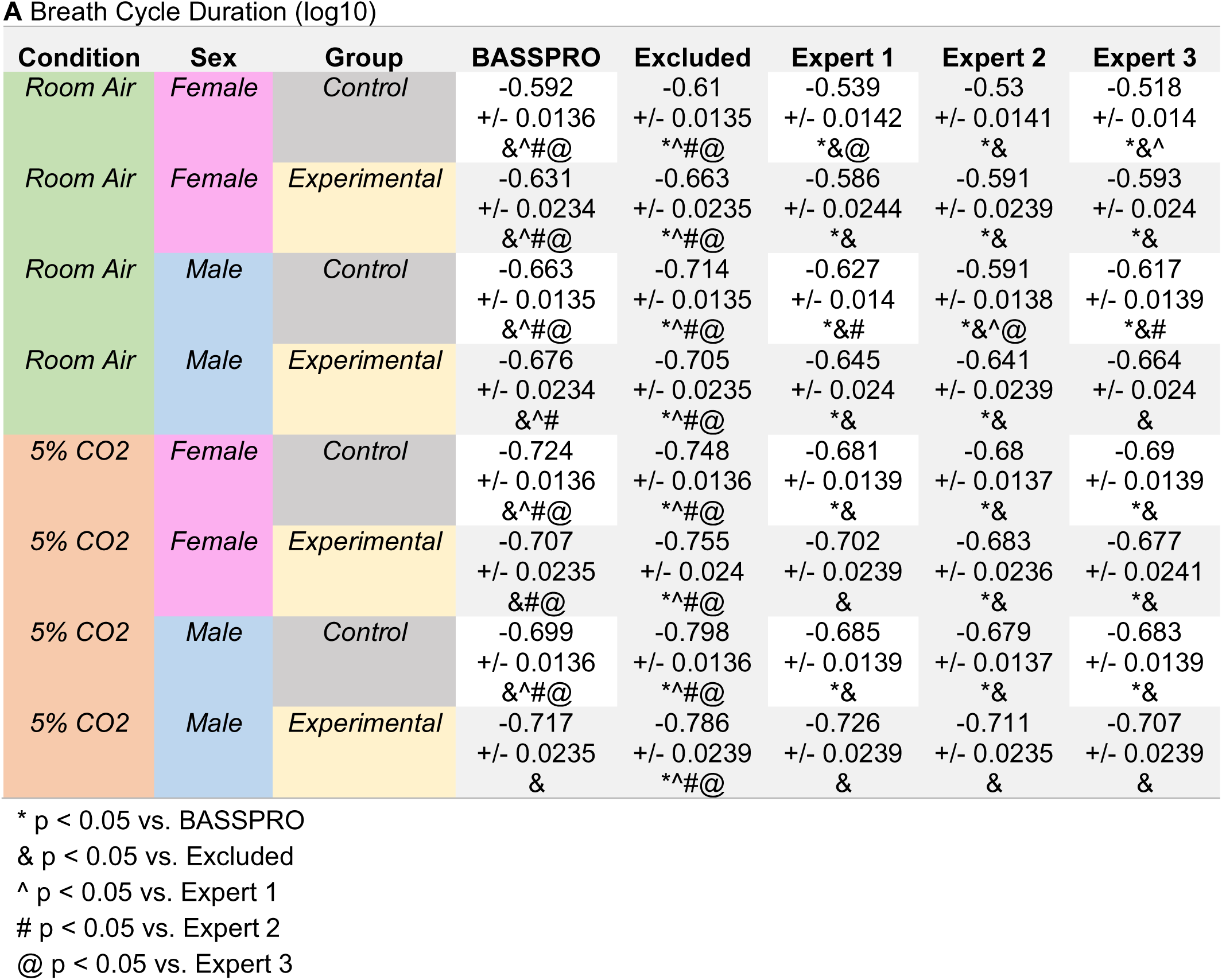
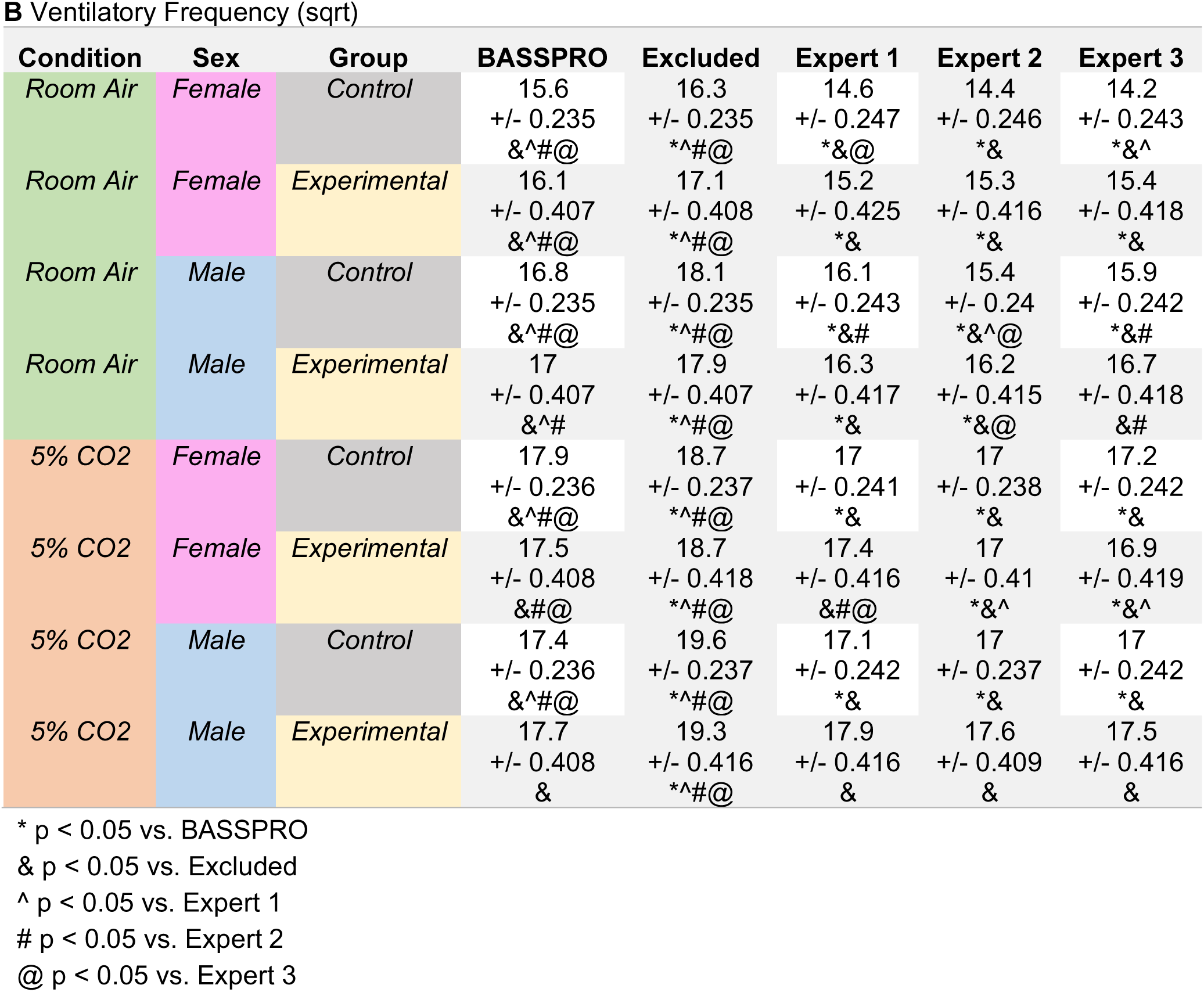
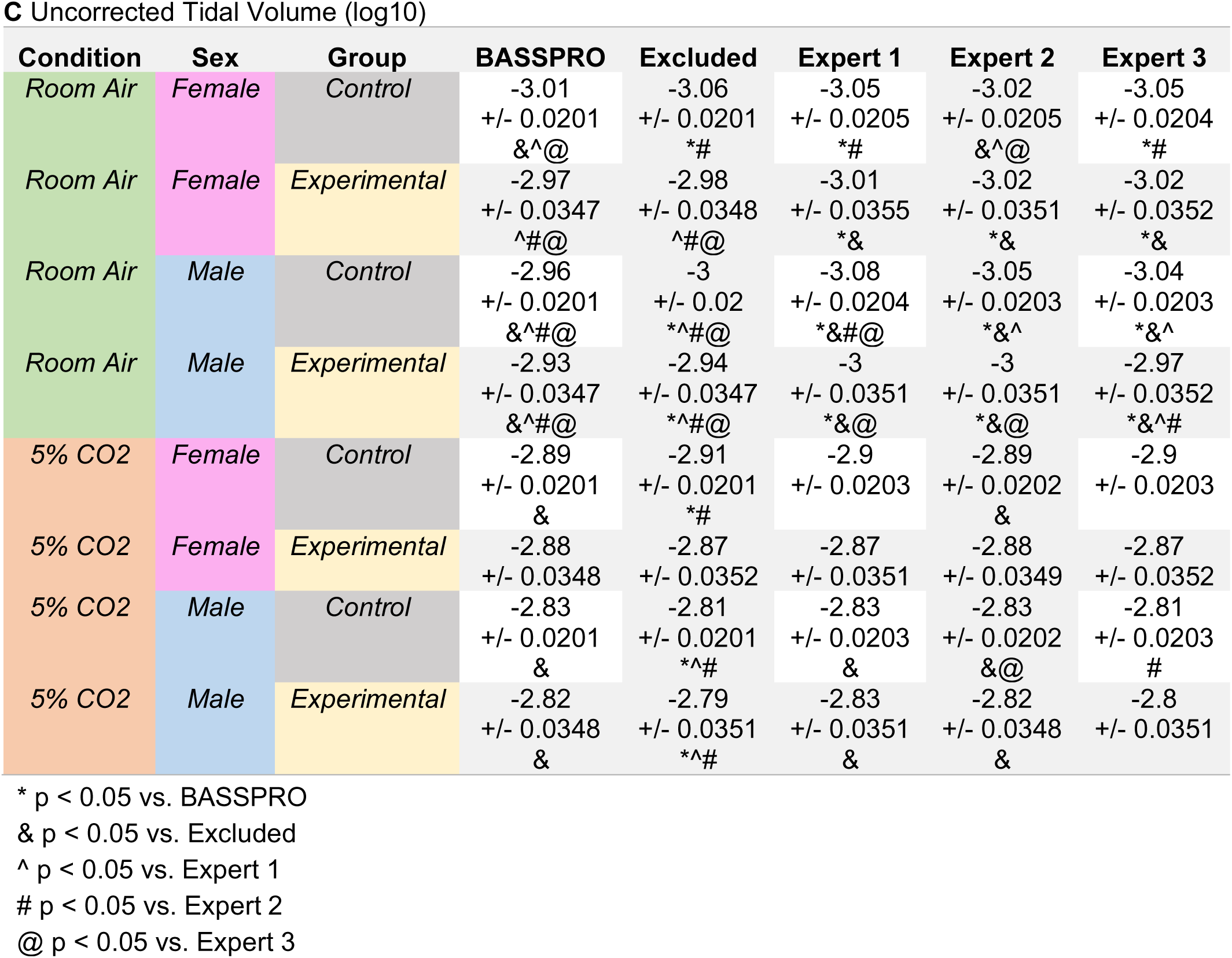
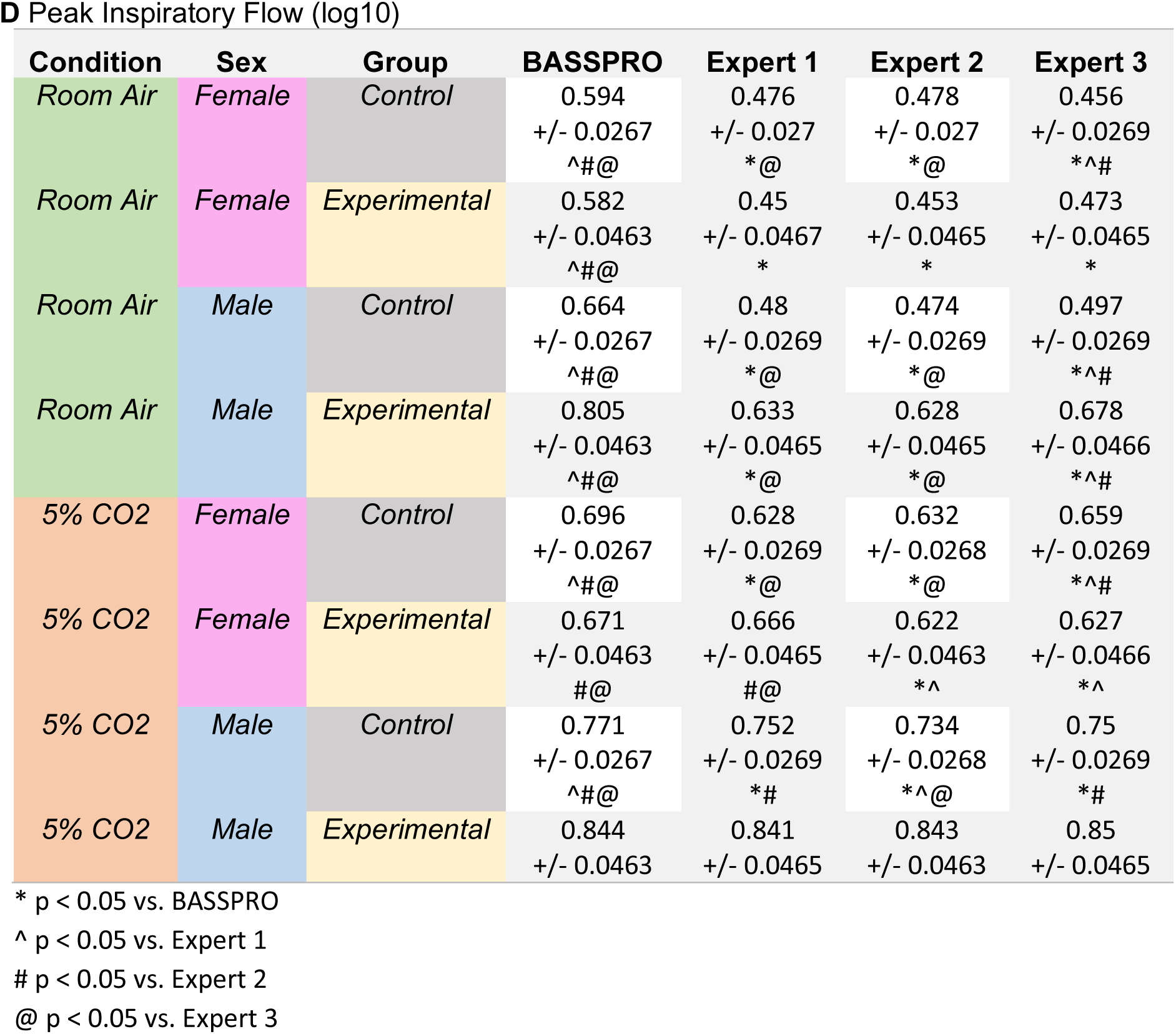
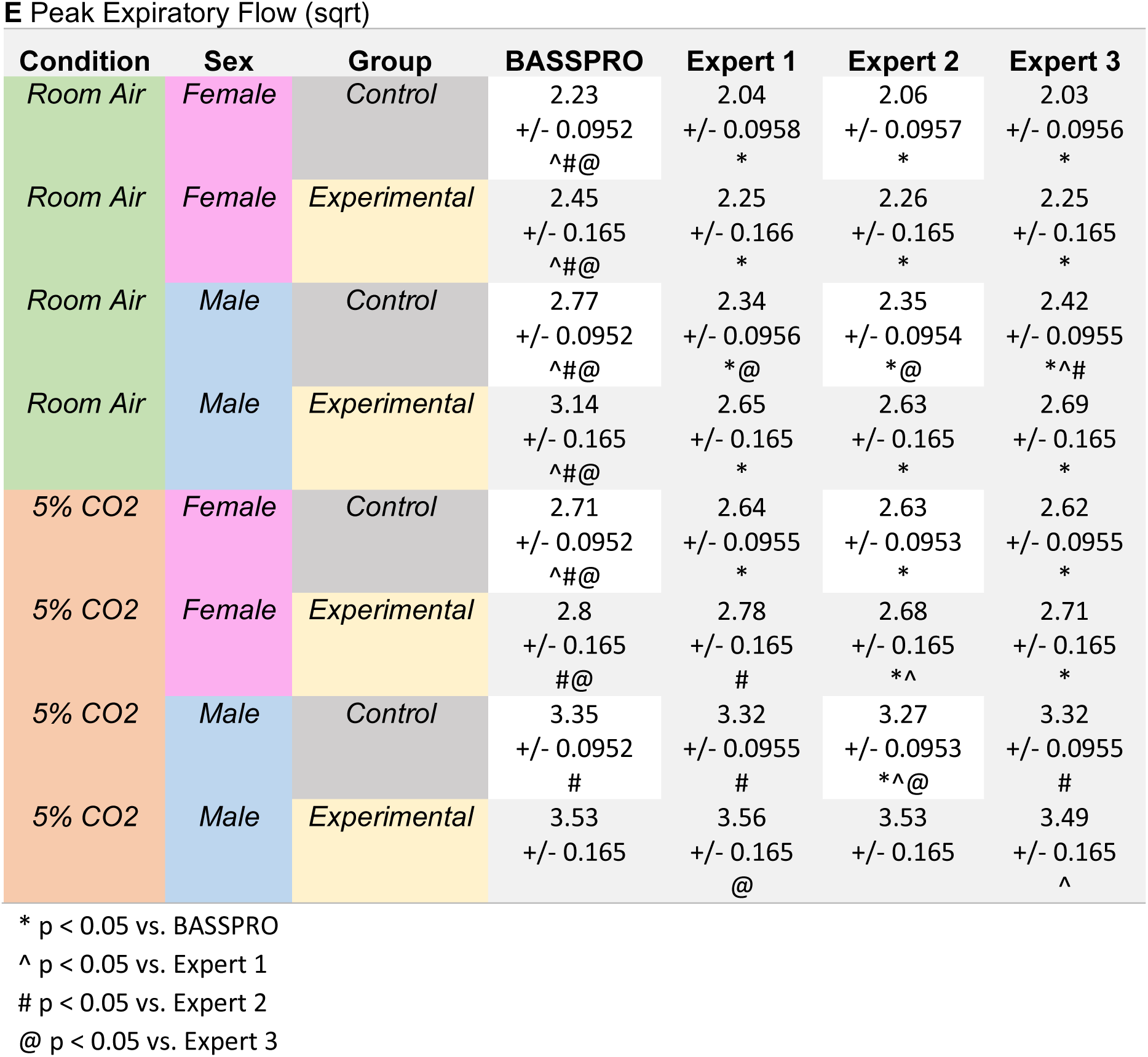
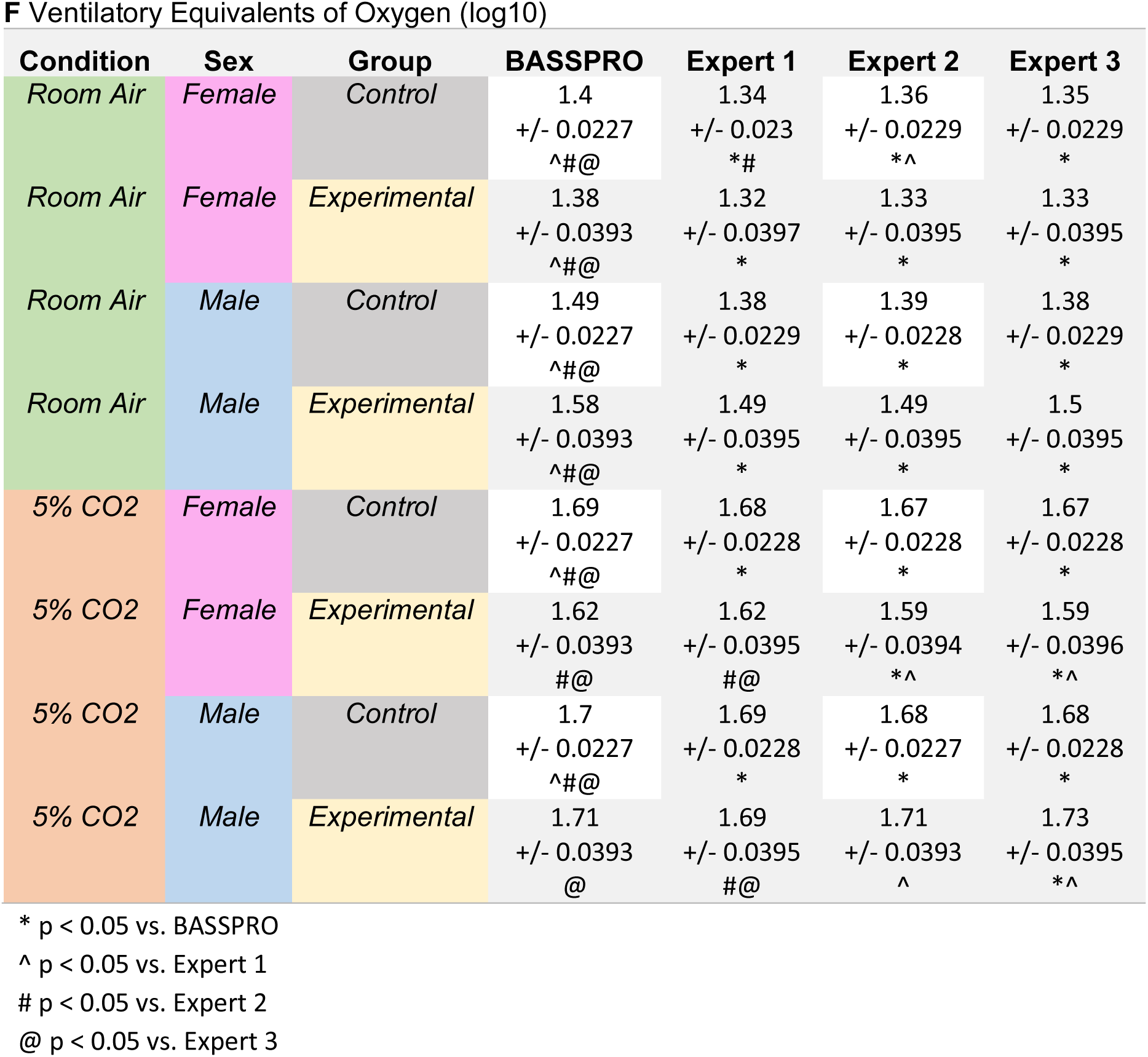
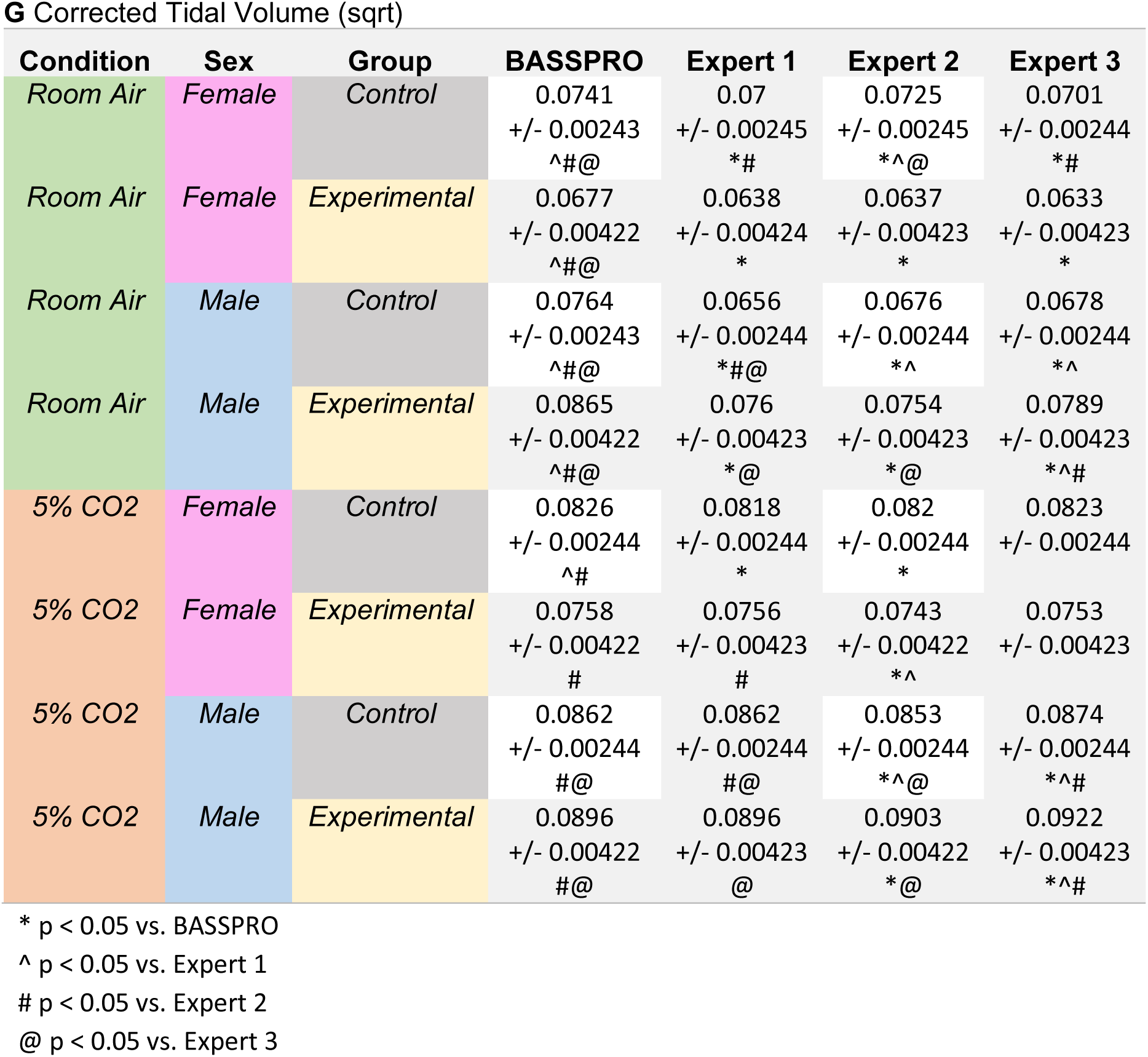
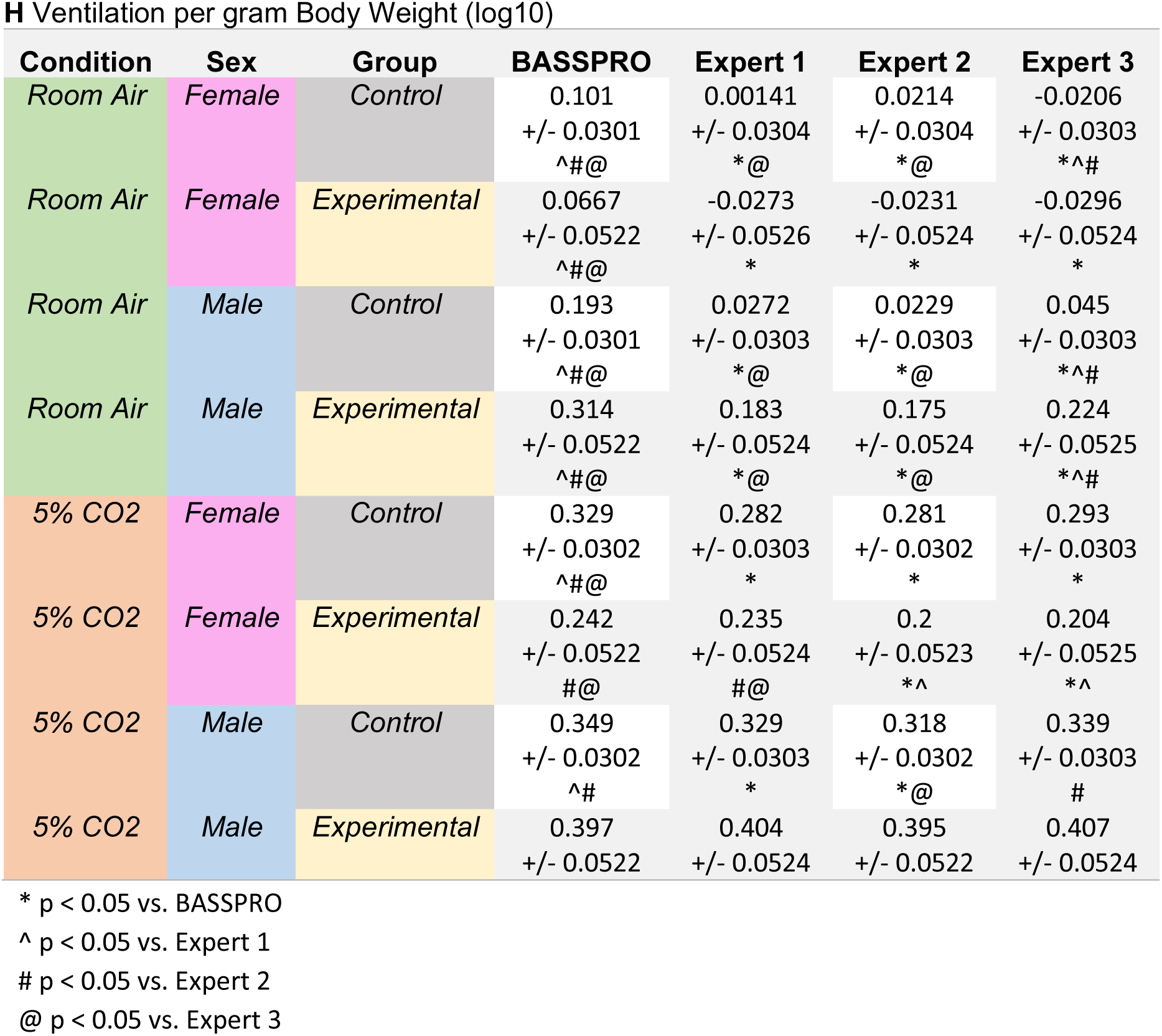
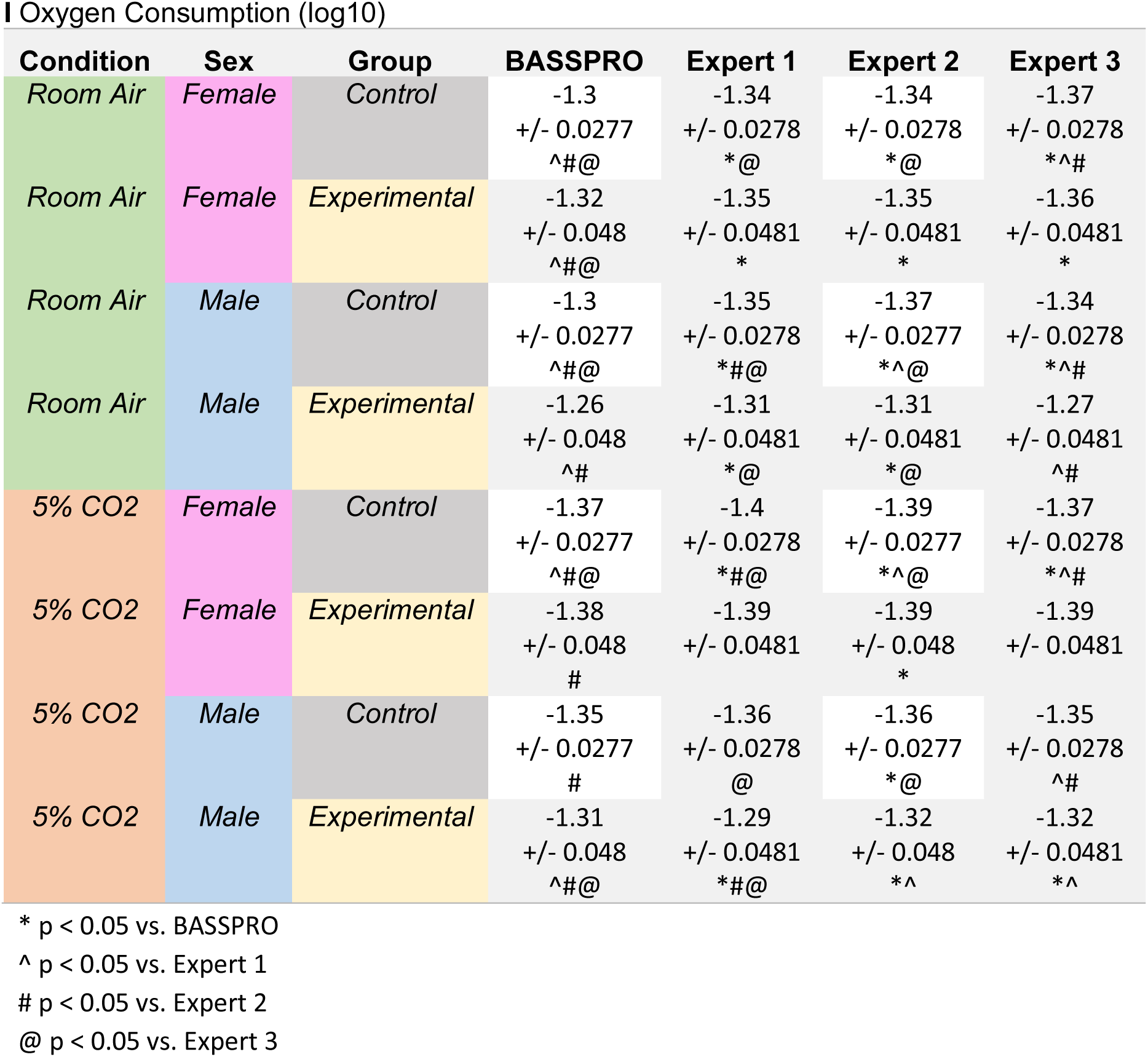
Average values for basic respiratory variables in validation study comparing BASSPRO automated breath selections to manual selections from 3 experts. Values present as mean ± standard error mean. Symbols following averages indicate significant differences between indicated groups. Reference group for identified significance are outlined below each table.

**Supplemental Table 4.**
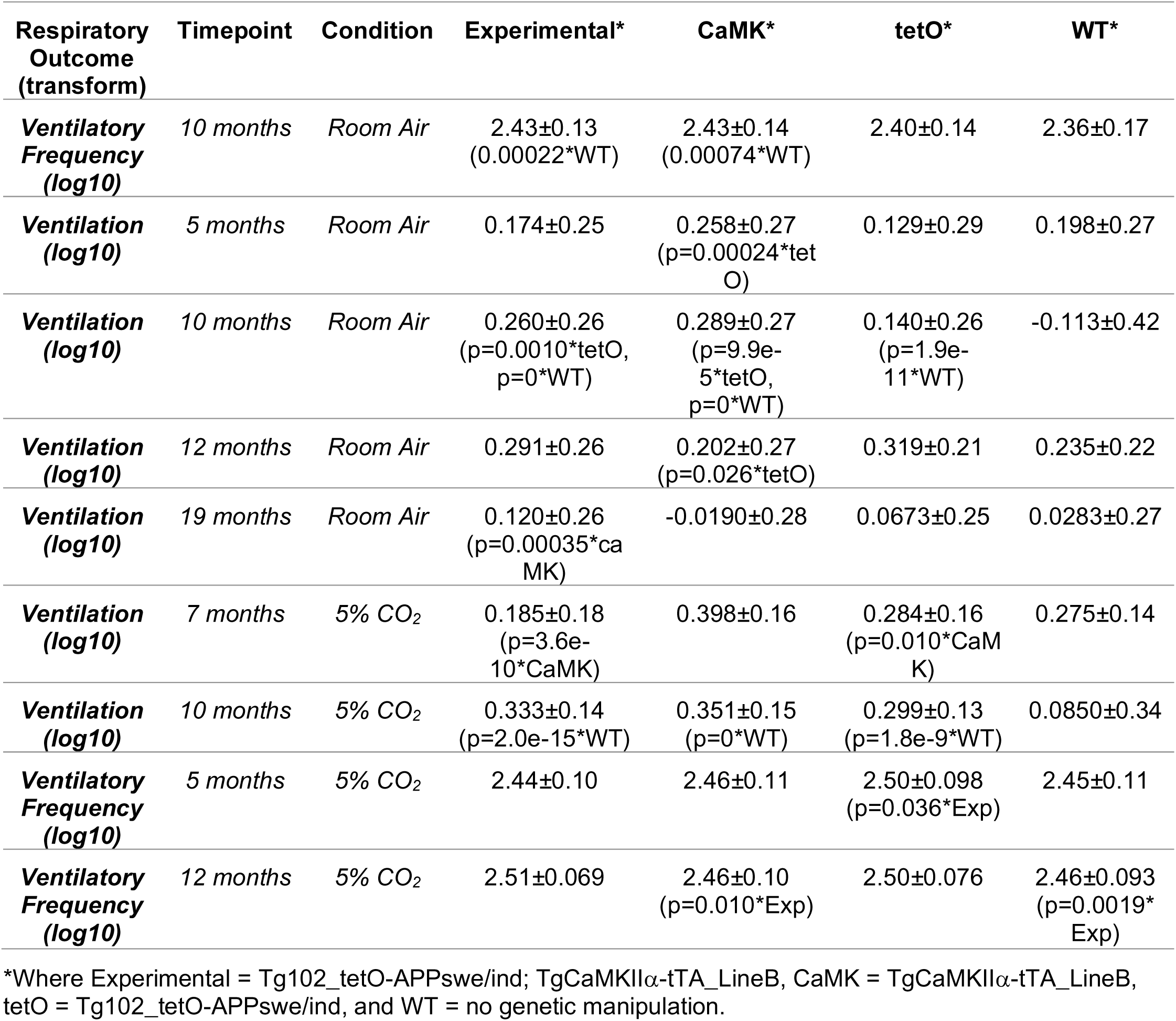
Average values and p-values for select ventilatory frequency and ventilation comparisons showing control variation in the Alzheimer’s dataset. *Presented as average ± SEM (p=p-value* Comparison Group).

| Respiratory Outcome (transform) | Timepoint | Condition | Experimental* | CaMK* | tetO* | WT* |
| --- | --- | --- | --- | --- | --- | --- |
| **Ventilatory Frequency (log10)** | 10 months | Room Air | 2.43±0.13<br>(0.00022*WT) | 2.43±0.14<br>(0.00074*WT) | 2.40±0.14 | 2.36±0.17 |
| **Ventilation (log10)** | 5 months | Room Air | 0.174±0.25 | 0.258±0.27<br>(p=0.00024*tetO) | 0.129±0.29 | 0.198±0.27 |
| **Ventilation (log10)** | 10 months | Room Air | 0.260±0.26<br>(p=0.0010*tetO, p=0*WT) | 0.289±0.27<br>(p=9.9e-5*tetO, p=0*WT) | 0.140±0.26<br>(p=1.9e-11*WT) | -0.113±0.42 |
| **Ventilation (log10)** | 12 months | Room Air | 0.291±0.26 | 0.202±0.27<br>(p=0.026*tetO) | 0.319±0.21 | 0.235±0.22 |
| **Ventilation (log10)** | 19 months | Room Air | 0.120±0.26<br>(p=0.00035*CaMK) | -0.0190±0.28 | 0.0673±0.25 | 0.0283±0.27 |
| **Ventilation (log10)** | 7 months | 5% CO<sub>2</sub> | 0.185±0.18<br>(p=3.6e-10*CaMK) | 0.398±0.16 | 0.284±0.16<br>(p=0.010*CaMK) | 0.275±0.14 |
| **Ventilation (log10)** | 10 months | 5% CO<sub>2</sub> | 0.333±0.14<br>(p=2.0e-15*WT) | 0.351±0.15<br>(p=0*WT) | 0.299±0.13<br>(p=1.8e-9*WT) | 0.0850±0.34 |
| **Ventilatory Frequency (log10)** | 5 months | 5% CO<sub>2</sub> | 2.44±0.10 | 2.46±0.11 | 2.50±0.098<br>(p=0.036*Exp) | 2.45±0.11 |
| **Ventilatory Frequency (log10)** | 12 months | 5% CO<sub>2</sub> | 2.51±0.069 | 2.46±0.10<br>(p=0.010*Exp) | 2.50±0.076 | 2.46±0.093<br>(p=0.0019*Exp) |

\*Where Experimental = Tg102\_tetO-APPswe/ind; TgCaMKII $\alpha$ -tTA\_LineB, CaMK = TgCaMKII $\alpha$ -tTA\_LineB, tetO = Tg102\_tetO-APPswe/ind, and WT = no genetic manipulation.

**Supplemental Table 5.** Number of mice included in analysis of hypercapnic (A) and hypoxic (B) ventilatory responses by age, gas condition, genotype, and sex. Average age (C) and weight (D) of all mice recorded by genotype, age group, hypercapnia or hypoxia exposure, and sex.

| Sex | Genotype | Condition | Age |  |  |  |  |  |  |
| --- | --- | --- | --- | --- | --- | --- | --- | --- | --- |
|  |  |  | 5 mo. | 7 mo. | 10 mo. | 12 mo. | 14 mo. | 16 mo. | 19 mo. |
| Female | CaM | 5% CO<sub>2</sub> | 6 | 6 | 6 | 6 | 6 | 5 | 6 |
|  |  | Room Air | 6 | 6 | 6 | 6 | 6 | 6 | 6 |
|  | Exp | 5% CO<sub>2</sub> | 6 | 6 | 6 | 5 | 6 | 5 | 6 |
|  |  | Room Air | 6 | 6 | 6 | 5 | 6 | 6 | 6 |
|  | tetO | 5% CO<sub>2</sub> | 4 | 4 | 4 | 2 | 4 | 4 | 3 |
|  |  | Room Air | 4 | 4 | 4 | 4 | 4 | 4 | 4 |
|  | WT | 5% CO<sub>2</sub> | 6 | 6 | 6 | 5 | 6 | 6 | 5 |
|  |  | Room Air | 6 | 6 | 6 | 5 | 6 | 6 | 6 |
| Male | CaM | 5% CO<sub>2</sub> | 6 | 6 | 6 | 5 | 6 | 5 | 5 |
|  |  | Room Air | 6 | 6 | 6 | 5 | 6 | 5 | 5 |
|  | Exp | 5% CO<sub>2</sub> | 5 | 5 | 5 | 3 | 5 | 5 | 5 |
|  |  | Room Air | 5 | 5 | 5 | 4 | 5 | 5 | 5 |
|  | tetO | 5% CO<sub>2</sub> | 6 | 6 | 6 | 6 | 6 | 5 | 5 |
|  |  | Room Air | 6 | 6 | 6 | 6 | 6 | 5 | 5 |
|  | WT | 5% CO<sub>2</sub> | 5 | 5 | 5 | 5 | 4 | 5 | 5 |
|  |  | Room Air | 5 | 5 | 5 | 5 | 4 | 5 | 5 |

**B** Number of mice included for analysis in hypoxia experiments by age group, gas condition, genotype, and sex.
| Sex | Genotype | Condition | Age |  |  |  |  |  |  |
| --- | --- | --- | --- | --- | --- | --- | --- | --- | --- |
|  |  |  | 5 mo. | 7 mo. | 10 mo. | 12 mo. | 14 mo. | 16 mo. | 19 mo. |
| Female | CaM | 10% O<sub>2</sub> | 4 | 3 | 4 | 4 | 4 | 4 | 3 |
|  |  | Room Air | 4 | 3 | 4 | 4 | 4 | 4 | 3 |
|  | Exp | 10% O<sub>2</sub> | 6 | 6 | 6 | 6 | 6 | 6 | 6 |
|  |  | Room Air | 6 | 6 | 6 | 6 | 6 | 6 | 6 |
|  | tetO | 10% O<sub>2</sub> | 6 | 6 | 6 | 6 | 6 | 6 | 6 |
|  |  | Room Air | 6 | 6 | 6 | 6 | 6 | 6 | 6 |
|  | WT | 10% O<sub>2</sub> | 6 | 6 | 6 | 6 | 3 | 6 | 5 |
|  |  | Room Air | 6 | 6 | 6 | 6 | 3 | 6 | 5 |
| Male | CaM | 10% O<sub>2</sub> | 5 | 6 | 6 | 6 | 6 | 5 | 6 |
|  |  | Room Air | 6 | 6 | 6 | 6 | 6 | 5 | 6 |
|  | Exp | 10% O<sub>2</sub> | 5 | 5 | 5 | 5 | 5 | 4 | 4 |
|  |  | Room Air | 5 | 5 | 5 | 5 | 5 | 5 | 5 |
|  | tetO | 10% O<sub>2</sub> | 5 | 6 | 6 | 6 | 6 | 5 | 5 |
|  |  | Room Air | 6 | 6 | 6 | 6 | 6 | 5 | 5 |
|  | WT | 10% O<sub>2</sub> | 5 | 5 | 5 | 5 | 5 | 5 | 5 |
|  |  | Room Air | 5 | 5 | 5 | 5 | 5 | 5 | 5 |

**C** Average age, in postnatal days, of mice by genotype, age group, hypercapnia or hypoxia exposure, and sex.
| Sex | Genotype | Condition | Age |  |  |  |  |  |  |
| --- | --- | --- | --- | --- | --- | --- | --- | --- | --- |
|  |  |  | 5 mo. | 7 mo. | 10 mo. | 12 mo. | 14 mo. | 16 mo. | 19 mo. |
| Female | CaM | Hypercapnia | 172.2 | 235.0 | 320.8 | 403.3 | 463.7 | 521.3 | 604.5 |
|  |  | Hypoxia | 176.2 | 241.0 | 322.3 | 412.0 | 468.0 | 525.0 | 607.0 |
|  | Exp | Hypercapnia | 171.8 | 235.7 | 320.7 | 406.0 | 464.0 | 520.7 | 603.0 |
|  |  | Hypoxia | 175.5 | 240.5 | 322.3 | 410.7 | 467.7 | 524.4 | 605.3 |
|  | tetO | Hypercapnia | 175.3 | 228.0 | 307.8 | 379.8 | 440.8 | 501.5 | 585.0 |
|  |  | Hypoxia | 180.3 | 230.5 | 312.0 | 388.5 | 444.5 | 504.5 | 588.0 |
|  | WT | Hypercapnia | 173.2 | 229.0 | 308.8 | 380.2 | 440.8 | 502.3 | 585.7 |
|  |  | Hypoxia | 178.2 | 231.7 | 313.2 | 387.7 | 446.7 | 505.3 | 588.6 |
| Male | CaM | Hypercapnia | 173.0 | 230.2 | 317.0 | 393.2 | 452.7 | 513.6 | 599.2 |
|  |  | Hypoxia | 175.8 | 236.0 | 319.7 | 398.7 | 462.7 | 517.4 | 602.2 |
|  | Exp | Hypercapnia | 169.6 | 233.0 | 317.0 | 390.3 | 450.4 | 512.2 | 597.8 |
|  |  | Hypoxia | 173.8 | 237.4 | 320.0 | 396.8 | 457.8 | 515.7 | 600.7 |
|  | tetO | Hypercapnia | 177.3 | 228.0 | 306.7 | 388.2 | 448.2 | 501.6 | 586.2 |
|  |  | Hypoxia | 182.4 | 231.2 | 308.3 | 396.5 | 455.8 | 506.4 | 587.8 |
|  | WT | Hypercapnia | 172.4 | 229.8 | 314.0 | 387.8 | 444.3 | 508.4 | 592.4 |
|  |  | Hypoxia | 176.4 | 234.0 | 316.2 | 394.2 | 457.2 | 511.6 | 594.6 |

**D** Average weight, in postnatal days, of mice by genotype, age group, hypercapnia or hypoxia exposure, and sex.
| Sex | Genotype | Condition | Age |  |  |  |  |  |  |
| --- | --- | --- | --- | --- | --- | --- | --- | --- | --- |
|  |  |  | 5 mo. | 7 mo. | 10 mo. | 12 mo. | 14 mo. | 16 mo. | 19 mo. |
| Female | CaM | Hypercapnia | 31.10 | 36.00 | 39.05 | 40.32 | 43.43 | 43.88 | 45.81 |
|  |  | Hypoxia | 31.56 | 36.28 | 39.46 | 41.92 | 43.67 | 43.74 | 45.54 |
|  | Exp | Hypercapnia | 33.35 | 37.68 | 41.76 | 41.98 | 46.12 | 45.40 | 47.97 |
|  |  | Hypoxia | 32.99 | 37.80 | 42.00 | 42.34 | 44.55 | 45.85 | 47.91 |
|  | tetO | Hypercapnia | 36.38 | 35.82 | 38.50 | 41.08 | 42.11 | 45.03 | 46.71 |
|  |  | Hypoxia | 35.84 | 36.79 | 39.47 | 41.73 | 42.55 | 44.98 | 45.88 |
|  | WT | Hypercapnia | 30.42 | 31.11 | 33.68 | 33.59 | 38.18 | 40.39 | 39.92 |
|  |  | Hypoxia | 29.79 | 31.92 | 34.74 | 35.53 | 36.59 | 39.47 | 39.58 |
| Male | CaM | Hypercapnia | 42.91 | 44.40 | 47.46 | 47.86 | 47.87 | 49.98 | 51.05 |
|  |  | Hypoxia | 41.34 | 43.00 | 46.74 | 48.15 | 48.63 | 49.52 | 50.25 |
|  | Exp | Hypercapnia | 40.30 | 44.24 | 44.16 | 47.07 | 47.70 | 48.76 | 48.43 |
|  |  | Hypoxia | 40.18 | 43.39 | 44.36 | 46.80 | 48.53 | 47.56 | 46.13 |
|  | tetO | Hypercapnia | 44.37 | 45.59 | 49.66 | 49.37 | 52.73 | 52.21 | 54.69 |
|  |  | Hypoxia | 43.15 | 45.10 | 48.94 | 50.02 | 52.67 | 51.91 | 54.07 |
|  | WT | Hypercapnia | 43.48 | 46.33 | 48.33 | 48.41 | 51.30 | 49.30 | 52.42 |
|  |  | Hypoxia | 41.94 | 45.10 | 48.01 | 48.99 | 50.11 | 49.71 | 52.57 |

**Supplemental Table 6.** All libraries and packages used by A) BASSPRO, B) STAGG, and C) the GUI and subGUIs. Citations are included for each library/ package.

| Library | Application | Citation |
| --- | --- | --- |
| argparse | This library collects arguments that are provided in the command line when executing the script so that they can be easily accessed as variables | 20 |
| csv | This library assists with reading in data from .csv files such as the metadata and auto_criteria settings files | 20 |
| datetime | This library provides a data structure for handling dates and time and applying mathematic operations to them – this is used as part of the logging system for tracking events through the execution of the script | 20 |
| logging | This library aids in generating contextualized logging feedback to the console as well as a log file. | 20 |
| math | This library provides additional functions for numerical operations and is currently used to aid in verifying if N/A values are present. | 20 |
| numpy | This library includes several classes and functions that are used including arrays for storing values as well as mathematic and filter generating functions | 21 |
| os | This library provides access to functions for file i/o and filepath creation and parsing | 20 |
| pandas | This library is used for its DataFrame class that is used to store several variables including the input signals and the detected breaths and their derived parameters. Several methods that act upon the DataFrame are utilized for filtering and applying calculations to the data. | 22 |
| re | This library is used for regular expression parsing of text strings and is used for parsing MUID and PlyUID information | 20 |
| scipy | This library provides functions used for signal processing including high and low pass filters that are used to reduce noise in the derived flow signal. Additionally, this library's *linregress* function is used for the creation of calibration curves (O<sub>2</sub> and CO<sub>2</sub> concentrations vs. voltages). | 23 |
| statistics | This library is used to provide access to statistical functions such as median | 20 |
| sys | This library is used to connect the systems stdout to logging functions, used for displaying logging messages in the console | 20 |
| tkinter | This library is used to provide a minimal GUI when running the BASSPRO script separately from the GUI | 20 |

**B** All packages used by STAGG.
| Package | Application | Citation |
| --- | --- | --- |
| argparser | Allows for operation of the R module, STAGG, from the command line terminal | 24 |
| data.table | Reformatting of data tables in preparation for manipulations | 25 |
| ggpubr | Graphing and data visualization | 26 |
| ggthemes | Additional settings and configurations for graphing | 27 |
| kableExtra | Builds common complex tables and manipulates table styles | 28 |
| lme4 | Fits linear and generalized linear mixed-effects models | 29 |
| lmerTest | Provides p-values for linear mixed-effects models | 30 |
| magrittr | Improves readability and intuitiveness of code | 31 |
| multcomp | Fits linear mixed-effects models and multiple comparisons | 32 |
| openxlsx | Reads, writes, and edits xlsx files | 33 |
| pandoc | Used for R markdown rendering | 34 |
| RColorBrewer | Generates colors for graphing | 35 |
| reshape2 | Used to transform data table formats for some optional graphs | 36 |
| rjson | Allows import of .json formatted files | 37 |
| rmarkdown | Produces html summary file with all .svg output files in one page | 38–40 |
| svglite | Produces user-friendly editable .svg image files. | 41 |
| tidyselect | Allows for verbiage consistency between packages | 42 |
| tidyverse | Data management and formatting; include dplyr and ggplot2 packages | 43 |
| xtable | Generates tables from functional outputs | 44 |

**C** All packages used by the GUI.
| Library | Application | Citation |
| --- | --- | --- |
| numpy | This library includes several classes and functions that are used including arrays for storing values as well as mathematic and filter generating functions | 21 |
| pandas | This library is used for its DataFrame class that is used to store several variables including the input signals and the detected breaths and their derived parameters. Several methods that act upon the DataFrame are utilized for filtering and applying calculations to the data. | 22 |
| pyodbc | This library provides methods and utilities for accessing ODBC databases in Python. | 45 |
| PyQt5 | This library is used for rendering graphics and GUI components like buttons and input boxes. It also manages processes that run parallel to the main GUI thread. | 46 |
| PyQt5-Qt5 | This library is a subset of a Qt installation that is required by PyQt5. | 46 |
| PyQt5-sip | This library provides Python bindings for C/C++ in support of PyQt5. | 47 |
| python-dateutil | This library extends the Python native package *datetime* with extra utilities for working with dates and times | 48 |
| pytz | This library provides Olson timezone calculations. | 49 |
| six | This library provides smooth compatibility between libraries written for both Python 2 and Python 3. | 50 |

**Supplemental Table 7.** Minimum and recommended machine requirements for running Breathe Easy.

|  | *Minimum Hardware* | *Recommended Hardware* |
| --- | --- | --- |
| *Operating System* | Windows 8 | Windows 10+ |
| *System Architecture* | 64-bit x86 | 64-bit x86 |
| *Diskspace* | Minimum 5 GB | Minimum 50 GB |
| *RAM* | Minimum 4 GB | Minimum 16 GB |

## References

1. Spinieli, R. L., Ben Musa, R., Kielhofner, J., Cornelius-Green, J. & Cummings, K. J. Orexin contributes to eupnea within a critical period of postnatal development. Am. J. Physiol. Regul. Integr. Comp. Physiol. 321, R558–R571 (2021).

2. Brown, A. G., Thapa, M., Hooker, J. W. & Ostrowski, T. D. Impaired chemoreflex correlates with decreased c-Fos in respiratory brainstem centers of the streptozotocin-induced Alzheimer’s disease rat model. Exp. Neurol. 311, 285–292 (2019).

3. Ebel, D. L., Torkilsen, C. G. & Ostrowski, T. D. Blunted Respiratory Responses in the Streptozotocin-Induced Alzheimer’s Disease Rat Model. J. Alzheimers Dis. 56, 1197– 1211 (2017).

4. Sun, J. J., Huang, T.-W., Neul, J. L. & Ray, R. S. Embryonic hindbrain patterning genes delineate distinct cardio-respiratory and metabolic homeostatic populations in the adult. Sci. Rep. 7, 9117 (2017).

5. Ward, C. S. et al. Loss of mecp2 function across several neuronal populations impairs breathing response to acute hypoxia. Front. Neurol. 11, 593554 (2020).

6. Lusk, S. et al. A CRISPR toolbox for generating intersectional genetic mouse models for functional, molecular, and anatomical circuit mapping. BMC Biol. (2022).

7. Liguori, C. et al. Sleep-disordered breathing and the risk of Alzheimer’s disease. Sleep Med. Rev. 55, 101375 (2021).

8. Liguori, C. et al. Orexinergic system dysregulation, sleep impairment, and cognitive decline in Alzheimer disease. JAMA Neurol. 71, 1498–1505 (2014).

9. Macheda, T. et al. Chronic Intermittent Hypoxia Induces Robust Astrogliosis in an Alzheimer’s Disease-Relevant Mouse Model. Neuroscience 398, 55–63 (2019).

10. Osorio, R. S. et al. Interaction between sleep-disordered breathing and apolipoprotein E genotype on cerebrospinal fluid biomarkers for Alzheimer’s disease in cognitively normal elderly individuals. Neurobiol. Aging 35, 1318–1324 (2014).

11. Ancoli-Israel, S. et al. Cognitive effects of treating obstructive sleep apnea in Alzheimer’s disease: a randomized controlled study. J. Am. Geriatr. Soc. 56, 2076–2081 (2008).

12. Troche, M. S., Huebner, I., Rosenbek, J. C., Okun, M. S. & Sapienza, C. M. Respiratory- swallowing coordination and swallowing safety in patients with Parkinson’s disease. Dysphagia 26, 218–224 (2011).

13. Yagi, N. et al. Inappropriate Timing of Swallow in the Respiratory Cycle Causes Breathing- Swallowing Discoordination. Front. Physiol. 8, 676 (2017).

14. Pitts, T. Airway protective mechanisms. Lung 192, 27–31 (2014).

15. Fuzik, J. et al. Integration of electrophysiological recordings with single-cell RNA-seq data identifies neuronal subtypes. Nat. Biotechnol. 34, 175–183 (2016).

16. Kalia, M. Dysphagia and aspiration pneumonia in patients with Alzheimer’s disease. Metab. Clin. Exp. 52, 36–38 (2003).

17. Won, J. H., Byun, S. J., Oh, B.-M., Park, S. J. & Seo, H. G. Risk and mortality of aspiration pneumonia in Parkinson’s disease: a nationwide database study. Sci. Rep. 11, 6597 (2021).

18. Wada, H. et al. Risk factors of aspiration pneumonia in Alzheimer’s disease patients. Gerontology 47, 271–276 (2001).

19. Suttrup, I. & Warnecke, T. Dysphagia in parkinson’s disease. Dysphagia 31, 24–32 (2016).

20. Jankowsky, J. L. et al. Persistent amyloidosis following suppression of Abeta production in a transgenic model of Alzheimer disease. PLoS Med. 2, e355 (2005).

21. Jankowsky, J. L. & Zheng, H. Practical considerations for choosing a mouse model of Alzheimer’s disease. Mol. Neurodegener. 12, 89 (2017).

22. Wilson, C. SWAN: Single Wave Analysis Notebook. (GitHub, 2016).

23. Sun, Y. et al. INSMA: An integrated system for multimodal data acquisition and analysis in the intensive care unit. J. Biomed. Inform. 106, 103434 (2020).

24. Hsieh, Y.-H. et al. Brainstem inflammation modulates the ventilatory pattern and its variability after acute lung injury in rodents. J. Physiol. (Lond.) 598, 2791–2811 (2020).

25. Sunshine, M. D. & Fuller, D. D. Automated classification of whole body plethysmography waveforms to quantify breathing patterns. Front. Physiol. 12, 690265 (2021).

26. Khurram, O. U., Gransee, H. M., Sieck, G. C. & Mantilla, C. B. Automated evaluation of respiratory signals to provide insight into respiratory drive. Respir. Physiol. Neurobiol. 300, 103872 (2022).

27. Schielzeth, H. et al. Robustness of linear mixed-effects models to violations of distributional assumptions. Methods Ecol. Evol. (2020). doi:10.1111/2041-210X.13434

28. Seiler, A. et al. Prevalence of sleep-disordered breathing after stroke and TIA: A meta- analysis. Neurology 92, e648–e654 (2019).

29. Koutsoukou, A. et al. Respiratory mechanics in brain injury: A review. World J. Crit. Care Med. 5, 65–73 (2016).

30. Dlouhy, B. J. et al. Breathing Inhibited When Seizures Spread to the Amygdala and upon Amygdala Stimulation. J. Neurosci. 35, 10281–10289 (2015).

31. Han, H. J. et al. Strain background influences neurotoxicity and behavioral abnormalities in mice expressing the tetracycline transactivator. J. Neurosci. 32, 10574–10586 (2012).

32. Janke, E. et al. Machine Learning to Classify the Emotional States of Mice from Respiration. SSRN Journal (2022). doi:10.2139/ssrn.4106834

33. Mayford, M. et al. Control of memory formation through regulated expression of a CaMKII transgene. Science 274, 1678–1683 (1996).

34. Ray, R. S. et al. Impaired respiratory and body temperature control upon acute serotonergic neuron inhibition. Science 333, 637–642 (2011).

35. Martinez, V. K. et al. Off-Target Effects of Clozapine-N-Oxide on the Chemosensory Reflex Are Masked by High Stress Levels. Front. Physiol. 10, 521 (2019).

36. Bartlett, D. & Tenney, S. M. Control of breathing in experimental anemia. Respir Physiol 10, 384–395 (1970).

37. Drorbaugh, J. E. & Fenn, W. O. A barometric method for measuring ventilation in newborn infants. Pediatrics 16, 81–87 (1955).

## References

1. Lusk, S. et al. A CRISPR toolbox for generating intersectional genetic mouse models for functional, molecular, and anatomical circuit mapping. BMC Biol. (2022).

2. Dauger, S., Nsegbe, E., Vardon, G., Gaultier, C. & Gallego, J. The effects of restraint on ventilatory responses to hypercapnia and hypoxia in adult mice. Respir Physiol 112, 215–225 (1998).

3. Martinez, V. K. et al. Off-Target Effects of Clozapine-N-Oxide on the Chemosensory Reflex Are Masked by High Stress Levels. Front. Physiol. 10, 521 (2019).

4. Tankersley, C. G., Haxhiu, M. A. & Gauda, E. B. Differential CO(2)-induced c-fos gene expression in the nucleus tractus solitarii of inbred mouse strains. J. Appl. Physiol. 92, 1277–1284 (2002).

5. Silva, T. de M. E., et al. Machine learning approaches reveal subtle differences in breathing and sleep fragmentation in Phox2b-derived astrocytes ablated mice. J. Neurophysiol. (2021). doi:10.1152/jn.00155.2020

6. Bassi, M. et al. Central leptin replacement enhances chemorespiratory responses in leptin-deficient mice independent of changes in body weight. Pflugers Arch. 464, 145–153 (2012).

7. Romer, S. H. et al. Accessory respiratory muscles enhance ventilation in ALS model mice and are activated by excitatory V2a neurons. Exp. Neurol. 287, 192–204 (2017).

8. Gestreau, C. et al. Task2 potassium channels set central respiratory CO2 and O2 sensitivity. Proc. Natl. Acad. Sci. USA 107, 2325–2330 (2010).

9. Huang, T.-W. et al. Progressive changes in a distributed neural circuit underlie breathing abnormalities in mice lacking mecp2. J. Neurosci. 36, 5572–5586 (2016).

10. Fechtner, L., El Ali, M., Sattar, A., Moore, M. & Strohl, K. P. Fentanyl effects on breath generation in C57BL/6J and A/J mouse strains. Respir. Physiol. Neurobiol. 215, 20–29 (2015).

11. Toward, M. A., Abdala, A. P., Knopp, S. J., Paton, J. F. R. & Bissonnette, J. M. Increasing brain serotonin corrects CO2 chemosensitivity in methyl-CpG-binding protein 2 (Mecp2)-deficient mice. Exp. Physiol. 98, 842–849 (2013).

12. Ward, C. S. et al. Loss of mecp2 function across several neuronal populations impairs breathing response to acute hypoxia. Front. Neurol. 11, 593554 (2020).

13. Sun, J. J., Huang, T.-W., Neul, J. L. & Ray, R. S. Embryonic hindbrain patterning genes delineate distinct cardio-respiratory and metabolic homeostatic populations in the adult. Sci. Rep. 7, 9117 (2017).

14. Hill, R., Dewey, W. L., Kelly, E. & Henderson, G. Oxycodone-induced tolerance to respiratory depression: reversal by ethanol, pregabalin and protein kinase C inhibition. Br. J. Pharmacol. 175, 2492–2503 (2018).

15. Vogelgesang, S., Niebert, M., Bischoff, A. M., Hülsmann, S. & Manzke, T. Persistent expression of serotonin receptor 5b alters breathing behavior in male mecp2 knockout mice. Front. Mol. Neurosci. 11, 28 (2018).

16. Ray, R. S. et al. Impaired respiratory and body temperature control upon acute serotonergic neuron inhibition. Science 333, 637–642 (2011).

17. Alhaddad, H. et al. Gender and strain contributions to the variability of buprenorphine-related respiratory toxicity in mice. Toxicology 305, 99–108 (2013).

18. Lewanowitsch, T., Miller, J. H. & Irvine, R. J. Reversal of morphine, methadone and heroin induced effects in mice by naloxone methiodide. Life Sci. 78, 682–688 (2006).

19. Goineau, S., Rompion, S., Guillaume, P. & Picard, S. Ventilatory function assessment in safety pharmacology: optimization of rodent studies using normocapnic or hypercapnic conditions. Toxicol. Appl. Pharmacol. 247, 191–197 (2010).

20. Van Rossum, G. & Drake, F. L. Python 3 Reference Manual. (CreateSpace, 2009).

21. Harris, C. R. et al. Array programming with NumPy. Nature 585, 357–362 (2020).

22. McKinney, W. Data Structures for Statistical Computing in Python. (2010).

23. Virtanen, P. et al. SciPy 1.0: fundamental algorithms for scientific computing in Python. Nat. Methods 17, 261–272 (2020).

24. Shih, D. J. H. argparser: Comman-Line Argument Parser. (R, 2021).

25. Dowle, M. & Srinivasan, A. *data.table: Extension of “data.frame.”* (R, 2021).

26. Kassambara, A. ggpubr: “ggplot2” Based Publication Ready Plots. (R, 2020).

27. Arnold, J. B. ggthemes: Extra Themes, Scales, and Geoms for “ggplot2.” (R, 2021).

28. Zhu, H. kableExtra: Construct Complex Table with “kable” and Pipe Syntax. (R, 2021).

29. Bates, D., Maechler, M., Bolker, B. & Walker, S. Fitting Linear Mixed-Effects Models Using lme4. J Stat Softw 67, 1–48 (2015).

30. Kuznetsova, A., Brockhoff, P. B. & Christensen, R. H. B. lmerTest Package: Tests in Linear Mixed Effects Models. Journal of Statistical Sofware 82, 1–26 (2017).

31. Bache, S. M. & Wickham, H. magrittr: A Forward-Pipe Operator for R. (R, 2020).

32. Hothorn, T., Bretz, F. & Westfall, P. Simultaneous Inference in General Parametric Models. Biom J 50, 346–363 (2008).

33. Schauberger, P. & Walker, A. openxlsx: Read, Write and Edit xlsx Files. (R, 2020).

34. Pandoc - About pandoc. at <https://pandoc.org/>

35. Neuwirth, E. RColorBrewer: ColorBrewer Palettes. (R, 2014).

36. Wickham, H. Reshaping Data with the reshape Package. J. Stat. Softw. 21, (2007).

37. Couture-Beil, A. rjson: JSON for R. (R, 2018).

38. Allaire, J. J. et al. rmarkdown: Dynamic Documents for R. (R, 2021).

39. Xie, Y., Dervieux, C. & Riederer, E. R Markdown Cookbook. (Chapman and Hall/ CRC, 2020).

40. Xie, Y., Allaire, J. J. & Grolemund, G. R Markdown: The Definitive Guide. (Chapman and Hall/ CRC, 2018).

41. Wickham, H., et al. svglite: An “SVG” Graphics Device. (r-lib.org, 2022).

42. Henry, L. & Wickham, H. tidyselect: Select from a Set of Strings. (R, 2020).

43. Wickham, H. et al. Welcome to the tidyverse. Journal of Open Source Software 4, 1686 (2019).

44. Dahl, D. B., Scott, D., Roosen, C., Magnusson, A. & Swinton, J. xtable: Export Tables to LaTeX or HTML. (R, 2019).

45. Kleehammer, M. pyodbc. (GitHub, Inc., 2020).

46. Riverbank Computing. PyQt. (Riverbank Computing, 2021).

47. Riverbank Computing. PyQt-sip. (Riverbank Computing, 1998). at <https://www.riverbankcomputing.com/software/sip>

48. Niemeyer, G., Pieviläinen, T., de Leeuw, Y. & Ganssle, P. python-dateutil. (GitHub, 2003). at <https://github.com/dateutil/dateutil>

49. Bishop, S. pytz. (PyPI, 2008).

50. Peterson, B. six. (PyPI, 2021). at <https://pypi.org/project/six/>

